# A closed feedback between tissue phase transitions and morphogen gradients drives patterning dynamics

**DOI:** 10.1101/2025.06.06.658228

**Authors:** Camilla Autorino, Diana Khoromskaia, Louise Harari, Elisa Floris, Harry Booth, Cristina Pallares-Cartes, Vesta Petrasiunaite, Michael Dorrity, Bernat Corominas-Murtra, Zena Hadjivasiliou, Nicoletta I. Petridou

## Abstract

During development mechanochemical cues in the cell microenvironment are translated into signalling to drive cell fate decisions. As cells differentiate collectively, it raises the question of how tissue-level properties affect instructive cues of decision-making. Here, we show that a tissue rigidity phase transition guides patterning by tuning the length-scales and time-scales of morphogen signalling. By combining rigidity percolation theory, reaction-diffusion modelling, quantitative imaging, optogenetics and single-cell transcriptomics in zebrafish, we uncover dynamical global tissue rigidity patterns that actively shape the Nodal morphogen gradient by restricting ligand dispersal and accelerating its signalling activity. In this self-generated mechanism, Nodal, besides driving meso-endoderm fate specification, increases cell-cell adhesion strength via regulating planar cell polarity genes. Once adhesion strength reaches a critical point, it triggers a rigidity transition which collapses tissue porosity. The abrupt tissue reorganisation negatively feeds back on Nodal signalling impacting both its length-scales, by limiting Nodal diffusivity, and its time-scales, by speeding up the expression of its antagonist Lefty, thereby ensuring timely signal termination and robust patterning. Overall, we reveal how emergent properties set the spatiotemporal dynamics of morphogen gradients, uncovering macroscopic mechanisms of pattern formation.

## MAIN TEXT

Embryo development is driven by processes spanning levels of biological organization. Accordingly, cell fate decisions, while guided by local mechanochemical signals generated at the level of the cell, may integrate information conveyed by properties arising at the supracellular, or collective-level, such as tissue rigidity, pressure or geometry ^1–4^. How the cell’s macro-environment impacts cell fate decisions is still elusive, largely due to the lack of approaches integrating tissue properties from a collective standpoint.

Recent work has grounded the shift between collective tissue states, e.g., solid-like and fluid-like, to theoretical frameworks of material phase transitions ^5–8^. During phase transitions, the tissue collective material state changes abruptly when a smoothly varying cell control parameter, like cell-cell adhesion or cell shape, crosses a specific but universal value, the *critical point* ^9–17^. In turn, tissue material properties are ultimately guided by the developmental programs of cell fate specification, such as morphogen signals ^13,18-21^. For instance, tissue rheological measurements in zebrafish embryos showed that Nodal signalling together with non-canonical Wnt signalling promote tissue solidification ^13,18^, whereas in the avian skin, FGF and BMP signalling promote tissue solidification and fluidization, respectively ^20^. Morphogen transport itself may be influenced by local changes in tissue architecture that defines the geometry of the extracellular space where morphogens diffuse or are up-taken by cells ^22–28^. However, if and how morphogen signalling and dynamic switches in tissue architecture are coordinated via feedback loops, and what are the functional implications of such an interplay remain open questions.

We explore these questions in the early zebrafish embryo where spatiotemporal variations in blastoderm viscosity coincide with its exit from pluripotency ^18^, with the latter being driven by the Nodal morphogen ^29^. At the blastula stage, the blastoderm is a multi-layered tissue composed of pluripotent cells surrounded by interstitial fluid (**Fig. 1a, b**, t 0 min). The first patterning event occurs at the onset of epiboly, when Nodal ligands, synthesised in the Yolk Syncytial Layer (YSL), are secreted in the blastoderm forming a gradient along the Animal-Vegetal (A-V) axis (**Fig. 1a-a’**) ^30–32^. When the marginal cells receive Nodal, Smad2/3 gets phosphorylated and enters the nucleus ^33,34^ to initiate transcription of meso-endodermal genes, Nodal itself, and its inhibitor Lefty ^32,35,36^ (**Fig. 1a’**), defining the specification region of the meso-endodermal layer (**Fig. 1a, b**, t 60 min). At the same stage, an abrupt tissue fluidization occurs in the central blastoderm with the specification zone exhibiting viscosity more than an order of magnitude higher than the still pluripotent blastoderm ^18^ (**Fig. 1b**, t 30-60 min). The changes in blastoderm viscosity can be characterized as a rigidity phase transition occurring at critical points in two control parameters, cell connectivity (**Fig. S1a, a’**) and cell-cell adhesion strength (**Fig. S1b, b’**) ^13,15^. Although the pluripotent and meso-endodermal regions exhibit slight differences in these control parameters, they exhibit dramatic differences in tissue-scale rigidity (**Fig. S1c, c’**). This raises the hypothesis that subtle changes in cell properties, potentially in response to the graded Nodal concentration, can significantly influence how collective tissue rigidity, in turn, shapes the Nodal gradient.

**Figure 1.**
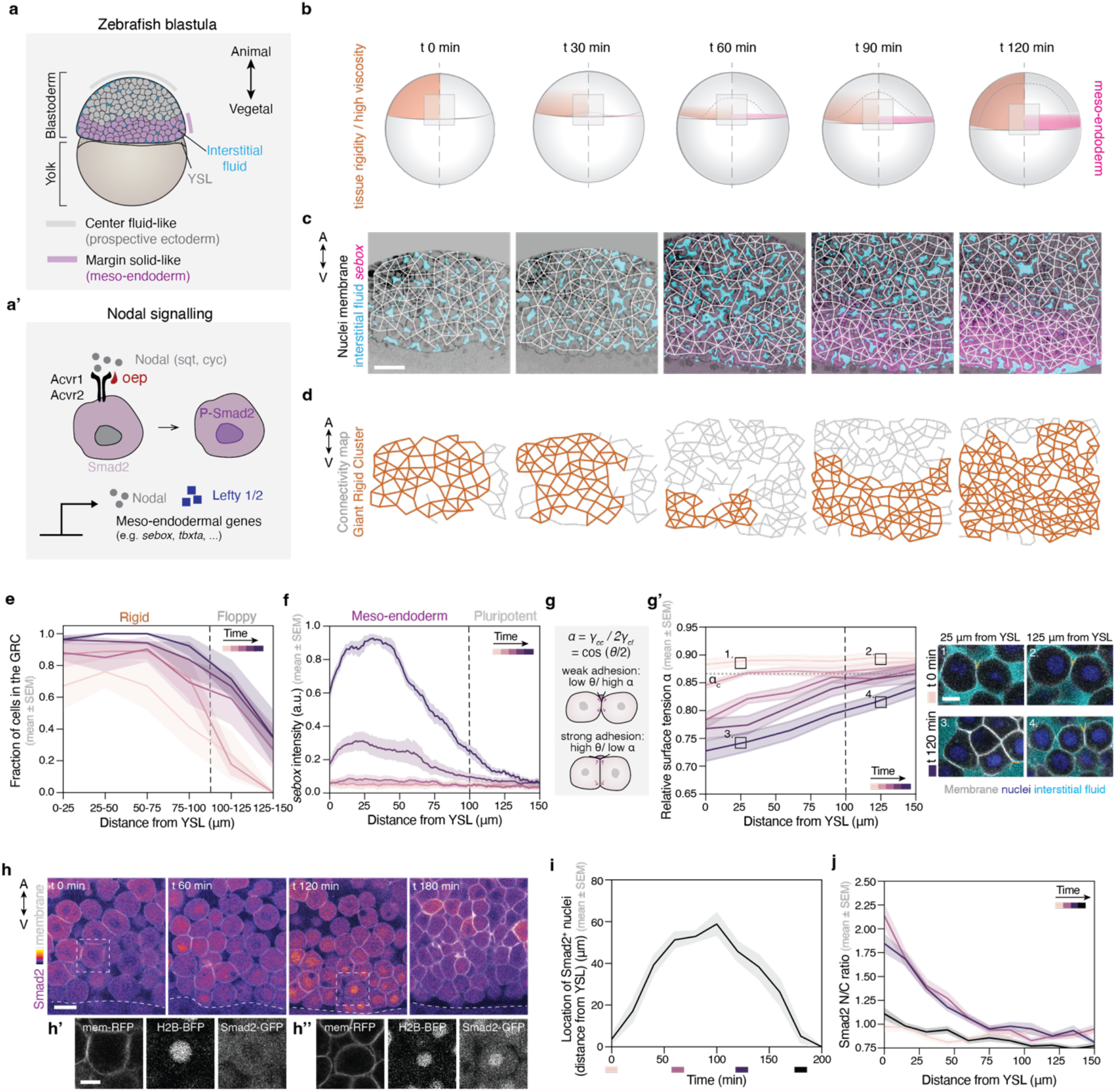
Meso-endodermal patterning in zebrafish correlates spatiotemporally with a tissue rigidity phase transition. **(a)** Schematic diagram of the zebrafish embryo at blastula stage, composed of the blastoderm, yolk and yolk syncytial layer, YSL. The blastoderm is composed of loosely-attached cells surrounded by interstitial fluid (cyan). The meso-endodermal layer is at the blastoderm margin. **(a’)** Schematic diagram of cell response to Nodal signalling including phosphorylation of Smad2/3, translocation to the nucleus and transcription of *nodal, lefty* and meso-endodermal genes. **(b)** Schematic diagram of the zebrafish embryo during meso-endoderm specification (right side), coinciding with changes in blastoderm rigidity and viscosity (left side). **(c)** Exemplary projected confocal sections from time lapse of transgenic embryos expressing eGFP in meso-endodermal progenitor cells (Tg(mezzo:eGFP), *sebox*) labelled with membrane-RFP, H2B-BFP and dextran-647 in the interstitial fluid, with overlaid connectivity maps. **(d)** Rigidity analysis of the cellular networks shown in (c), with the GRC shown in orange. **(e-f)** Plots of the distribution of the GRC (e) and *sebox* intensity (f), as a function of the distance from the YSL, over time, time intervals of 30 min. Dashed lines indicate the rigid-floppy boundary in (e) and the meso-endoderm - prospective ectoderm boundary in (f) (n = 4 embryos for (e) and n = 8 embryos for (f)). **(g)** Schematic diagram of the relative surface tension *α* as defined by the Young-Dupré relation, measured by the contact angle. **(g’)** Plot of the distribution of *α* as a function of the distance from the YSL, over time (left), time intervals of 30 min, and exemplary confocal images of the contact angles (right) in the different blastoderm regions and time points (regions 1, 3: cells close to YSL; regions 2, 4: cells further away from YSL) (n = 8 embryos). **(h)** Exemplary 2D confocal sections from live embryos labelled for membrane-RFP, H2B-BFP (only in h’ and h’’) and Smad2-GFP. The dotted line indicates the YSL. Dotted boxes indicate close-ups at time points t 0 min (h’) and at t 120 min (h’’). **(i)** Plot of the distribution of nuclear Smad2-positive cells across the A-V axis as a function of time. Shaded boxes indicate the time points shown in (h) (n = 6 embryos). **(j)** Plot of the Smad2-GFP N/C ratio intensity over time as a function of the distance from YSL (n = 3 embryos), time intervals of 60 min. A-V, Animal-Vegetal axis; GRC, Giant Rigid Cluster; N/C, Nuclear-to-cytoplasmic ratio; YSL, Yolk Syncytial Layer. Scale bars: 50 μm (c), 10 μm (g’, h’, h’’) 20 μm (h)

To identify a potential feedback regulation between morphogen gradients and emergent tissue properties, we here combined quantitative live imaging to map the tissue material state and Nodal morphogen signalling dynamics, together with genetics and optogenetics to modulate signalling and tissue phase transitions, respectively. This allowed us to mechanistically describe a closed feedback between Nodal gradient formation and tissue phase transitions. Together with a theoretical framework integrating tissue phase transitions with reaction-diffusion biochemical networks, we show that this feedback is crucial for tuning the Nodal length-scales and time-scales needed for robust patterning. Therefore, this work uncovers macroscopic mechanisms of morphogen gradient formation, where dynamic switches in collective tissue physical states do not simply define tissue deformability ^13,16,18,37–41^, but directly regulate the instructive cues setting the spatiotemporal dynamics of cell fate decisions.

### Patterning during a rigidity transition

To explore the interplay between meso-endodermal patterning and tissue material phase transitions, we quantified the joint spatiotemporal dynamics of cell fate and tissue material properties. As a readout of cell fate specification, we live imaged the ventrolateral marginal blastoderm of zebrafish embryos labelled for membrane, nuclei, interstitial fluid and the meso-endodermal marker *sebox* ^42^, starting at pluripotent stages (t 0 min) until the onset of gastrulation (t 120 min) (**Fig. 1b, c, f**). To quantify the material state of the blastoderm, we used rigidity percolation theory, a framework mapping the network representation of a material to its deformability ^43,44^. Embryonic tissues can be abstracted as cellular networks, in which cell centroids are nodes and cell-cell contacts are viscoelastic bonds ^13,15,45^ (**Fig. 1c**). Within this framework, the material response of the network can be inferred by evaluating the size of the largest network cluster within which nodes have no independent movements, a collective property of the network referred to as the Giant Rigid Cluster (GRC) (**Fig. 1d**, orange cluster). A transition in the GRC size occurs once the network crosses a critical point in the control parameter average cell-cell connectivity, <*k*>, defined as the average number of contacts per cell (*k*_*c*_ ≈ 4, ~⅔ of maximum connectivity) (**Fig. S1a, a’**) ^43,44,46^. Using this approach, we detect a transient spatial correlation between the specification zone and the rigid domain, where the former is secluded within the GRC at the blastoderm margin (**Fig. 1c-f**, t 60-90 min). This effectively creates a physical boundary between the specifying domain and the overlaying pluripotent tissue (**Fig. 1e, f**, dashed lines, **Fig. S1d, Movie 1**). The co-occurring changes in the two macroscopic properties, meso-endodermal fate and tissue rigidity, prompted us to ask if their microscale regulators are spatiotemporally coordinated.

We explored the above question by quantifying the dynamics of the microscopic components regulating meso-endodermal fate and tissue rigidity. For meso-endodermal fate, we focused on the Nodal gradient. We performed live imaging of Smad2 and quantified the number and spatial distribution of cells with nuclear Smad2 (**Fig. 1h, i, Fig. S1e, Movie 2**). In agreement with previous reports ^47–51^, we observe a short Nodal signalling range along the A-V axis, where only the first 3-4 most marginal cell tiers receive Nodal (closer to YSL), while cells retain the signal for ~3 hours (**Fig. 1i, Fig. S1e**). This pattern further matches Nodal activation levels as evaluated via live imaging and quantification of the nuclear-to-cytoplasmic (N/C) ratio signal (**Fig. 1j**) and P-Smad2/3 immunostaining (**Fig. S1f, g**), where a short-range signalling gradient is observed which steepens over time and eventually turns off.

We then quantified connectivity and GRC size along the A-V axis, in order to elucidate if the spatial rigidity pattern emerges as a function of spatial gradients in connectivity. We found that when the marginal tissue is still at pluripotency, the tissue is poised close to the critical point in connectivity, *k*_*c*_ (**Fig. S1h**). As specification progresses, connectivity increases and crosses the critical point in the most marginal cells (**Fig. S1h**). The changes in connectivity seem to mainly result from a gradient in cell-cell adhesion strength (**Fig. 1g, g’**), which also acts as a control parameter of tissue rigidity (**Fig. S1b**) ^15^. Cell-cell adhesion strength can be inferred using the Young-Dupré relation, where a non-dimensional parameter *α*, defined as the ratio between cell-cell and cell-fluid surface tensions acting at the contact, can be calculated from the angle formed at the contact edge (**Fig. 1g, g’, Fig. S1b’**) ^13,15,52-56^. Quantifying *α* along the A-V axis revealed that similarly to connectivity, at pluripotency, the tissue is poised close to the critical point of the relative surface tension (*α*_*c*_ ≈ 0.8662) (**Fig. 1g’**, box 1, **Fig. S1i**), and during specification, the most marginal cells reduce their *α* values below *α*_*c*_ and become more adhesive (**Fig. 1g’**, box 3, **Fig. S1i**). Given that the critical points in *α* and <*k*> are crossed along the A-V axis, it suggests that during the rigidity transition, the initial spatial isotropy of the tissue is broken, polarizing the GRC towards the specification zone. To test if the slight gradient in the control parameters is sufficient to polarize tissue rigidity, we simulated networks and tissues with and without a gradient in <*k*> and *α*, respectively, and explored potential changes in the GRC location along the A-V axis (**Fig. S1j-m**). Networks with spatially isotropic connectivity values close to criticality position the GRC at any location along the axis (**Fig. S1j, k**). In contrast, under the same <*k*> values, a slight gradient in connectivity breaks the symmetry and polarizes the GRC towards the super-critical connectivity regions (**Fig. S1j, k**, Supplementary Theory Note). The same trends are seen for *α*, where *in silico* rigidification in the presence of a slight gradient in *α* is sufficient to polarize the GRC along the A-V axis (**Fig. S1l, m**, Supplementary Theory Note).

Overall, the above results show that during patterning, the Nodal morphogen gradient arises together with a cell-cell adhesion gradient. The latter triggers a symmetry breaking-like event in the initially isotropic tissue material state, correlating the spatiotemporal dynamics of tissue rigidification to specification.

### A Nodal - tissue rigidity feedback loop

The correlation between the spatiotemporal dynamics of patterning and tissue rigidity prompted us to ask whether Nodal signalling and the control parameters that drive tissue rigidification are functionally linked. We first explored if Nodal signalling drives tissue rigidification. To this end, we quantified *α*, connectivity and GRC in embryos that either lack Nodal signalling (*MZoep*, mutant for Nodal co-receptor ^33,57^) or have increased Nodal signalling (*MZlefty1/2*, mutant for Nodal inhibitors ^51^) (**Fig. 1a’**). In contrast to *wildtype* embryos, nuclear Smad2 signal is absent in *MZoep* embryos, confirming that Nodal signalling is impeded. Inversely, nuclear Smad2 region expands in *MZlefty1/2* embryos confirming that Nodal signalling is enhanced (**Fig. 2d, e, Fig. S2g, Movie 2**). In *MZoep* embryos, connectivity and *α* values remain at criticality (**Fig. 2b, c, Fig. S2a**) resulting in a random distribution of the GRC across the A-V axis (**Fig. 2a, a’, f, Fig. S2b**). This resembles the rigidity profiles of the *in silico* tissues without a gradient in their control parameter (**Fig. S1j-m**). Conversely, in *MZlefty1/2* mutants, connectivity and *α* values change faster (**Fig. 2b, c, Fig. S2a**), percolating tissue rigidity throughout the entire tissue (**Fig. 2a, a’, f, Fig. S2b**). Altogether, these experiments show that Nodal signalling sets the spatial rigidity pattern of the zebrafish margin by regulating cell-cell adhesion and polarizing a highly connected rigid cluster of cells towards the margin (**Fig. 2f, g**).

**Figure 2:**
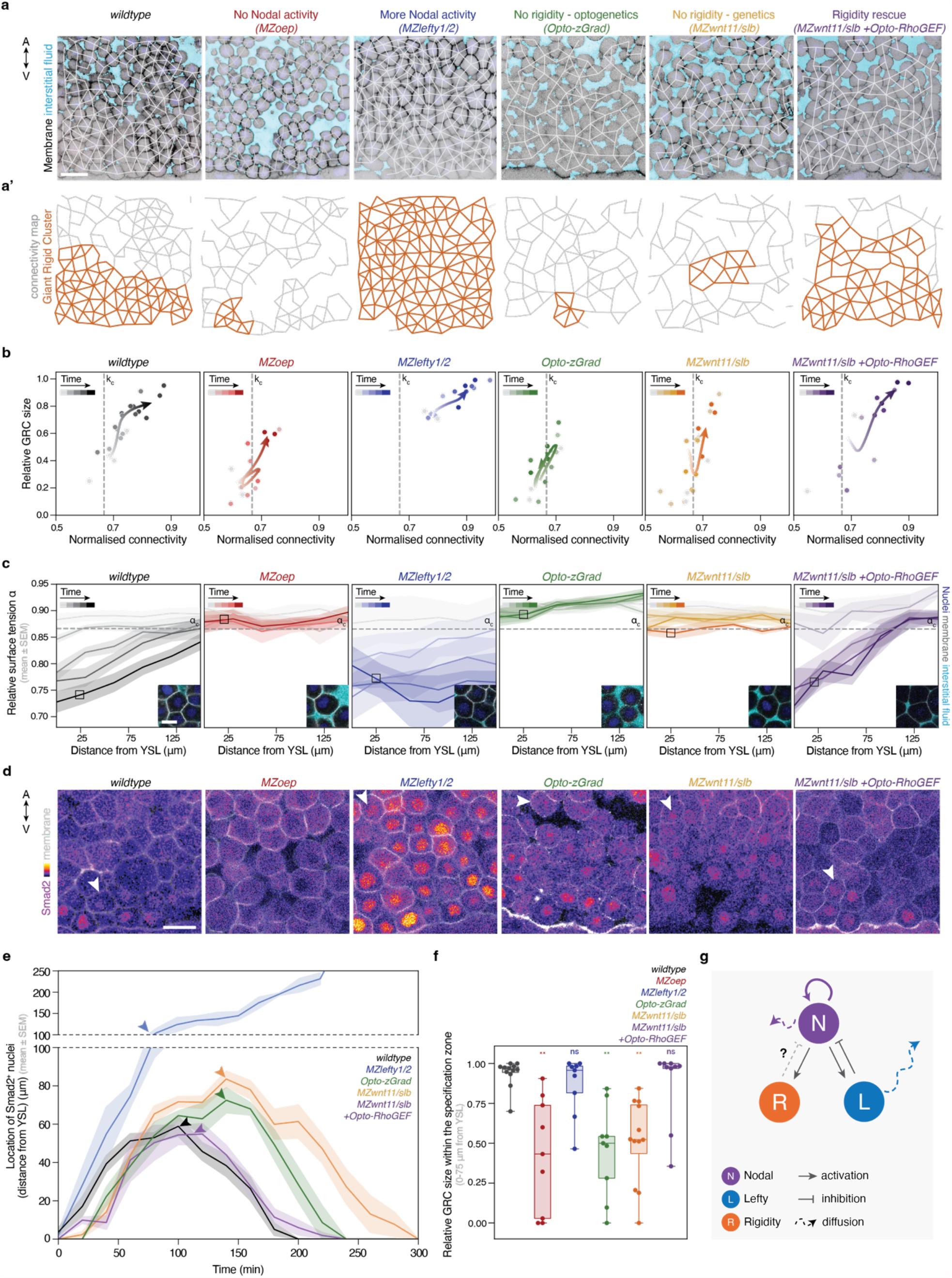
Nodal triggers a tissue rigidity phase transition along the A-V axis, which feedbacks to terminate Nodal signalling. **(a)** Exemplary 2D confocal sections at t 90 min for *wildtype, MZoep, MZlefty1/2, Opto-zGrad, MZwnt11/slb* and *MZwnt11/slb +Opto-RhoGEF* embryos labelled with membrane-RFP (α-catenin-Citrine in Opto-zGrad) and dextran-647 for interstitial fluid, with overlaid connectivity maps and their corresponding rigidity profiles (a’). **(b)** Plots of GRC size as a function of connectivity over time for the conditions shown in (a) (n = 4 embryos *wildtype*, n = 3 *MZoep*, n = 3 *MZlefty1/2*, n = 3 *Opto-zGrad*, n = 4 *MZwnt11/slb*, n = 3 *MZwnt11/slb +Opto-RhoGEF*). Dashed lines indicate *k*_*c*_. The arrow connects the mean connectivity and GRC values for each time point. **(c)** Plots of the relative surface tension *α* as a function of the distance from the YSL over time for the conditions shown in (a) with exemplary high magnification confocal sections depicting the contact angles. Dashed lines indicate *α*_*c*_ (*wildtype*: n = 12274 contact angles, N = 8 embryos; *MZoep*: n = 4003, N = 3; *MZlefty1/2*: n = 3651, N = 3; *Opto-zGrad*: n = 2575, N = 3; *MZwnt11/slb*: n = 6224, N = 6; *MZwnt11/slb +Opto-RhoGEF*: n = 2386, N = 3). **(d)** Exemplary 2D confocal sections of embryos from the conditions described in (a) labelled with membrane-RFP and Smad2-GFP. The white arrowheads indicate the largest distance from the YSL with Smad2 positive nuclei. **(e)** Plot of the distribution of nuclear Smad2-positive cells as a function of time for the conditions shown in (a). Arrowheads indicate the time points at which the images in (d) were selected, corresponding to the peak of Nodal activity length-scale (n = 6 embryos *wildtype*, n = 3 *MZlefty1/2*, n = 3 *Opto-zGrad*, n = 4 *MZwnt11/slb*, n = 3 *MZwnt11/slb +Opto-RhoGEF*). **(f)** Quantification of GRC relative size in the specification zone (0-75 μm form YSL), for all the conditions in (a), from t 60 min to t 120 min. (n = 12 *wildtype*, n = 9 *MZoep*, n = 9 *MZlefty1/2*, n = 9 *Opto-zGrad*, n = 12 *MZwnt11/slb*, n = 9 *MZwnt11/slb +Opto-RhoGEF*) **(g)** Illustration of the Nodal-Lefty-Rigidity interaction network. The time axes shown in (b) and (c) correspond to time intervals of 30 min. A-V, Animal-Vegetal axis; GRC, Giant Rigid Cluster; YSL, Yolk Syncytial Layer. Statistics: (f) ANOVA and Kruskal-Wallis multiple comparisons test, compared to *wildtype*. (*MZoep* p-value = 0.002. *MZlefty1/2* p-value = 0.99, *Opto-zGrad* p-value = 0.002, *MZwnt11/slb* p-value = 0.002, *MZwnt11/slb +Opto-RhoGEF* p-value = 0.99). Scale bars: 25 μm (a), 10 μm (c), 20 μm (d)

As a next step, we asked if the collective material state affects meso-endoderm specification. To address this, we monitored Nodal dynamics using Smad2 live imaging in tissues with altered material properties. Our previous findings showed that tissue rigidity can be precisely altered by finetuning the relative surface tension *α* using optogenetic tools ^15^. To this end, we used a degradation-based optogenetic system (*Opto-zGrad*) ^15,58,59^ (**Fig. S2c**) to maintain *α* close to *α*_*c*_ and thus, inhibit the polarisation of the rigid domain. Specifically, we mildly reduced α-catenin levels by inducing its degradation in a spatiotemporally resolved manner (**Fig. S2d, d’**) to inhibit the decrease of *α* in the most marginal cells (**Fig. 2a-c, f, Fig. S2a, b**). Remarkably, when examining Nodal signalling we observed that more cells, and cells further away from the YSL, display nuclear Smad2 indicating a shift in the length-scales of Nodal signalling in the fluidised embryos (**Fig. 2d, e, Fig. S2g**). Moreover, Smad2 remains active for a longer period, suggesting that the tissue material state also influences the time-scale of Nodal signalling (**Fig. 2e, Fig. S2g**). Importantly, the change in Smad2 dynamics is not due to indirect effects of cell-cell adhesion on other signalling pathways, such as Wnt/β-catenin ^60^, as there is no difference in β-catenin staining between *wildtype* and *Opto-zGrad* embryos (**Fig. S2f-f’’**). Taken together, these results show that tissue rigidity can impact Nodal signalling by restricting its spatial and temporal dynamics.

Given that the adhesion gradient and emergent rigidity observed in *wildtype* embryos are instructed by Nodal, we hypothesized that Nodal controls its own termination not only biochemically - via the activation of its inhibitor Lefty-but also mechanically, by regulating tissue rigidity. To explore such potential negative feedback, we first asked how Nodal might influence cell-cell adhesion strength. Previous work has shown that the expression of the planar cell polarity (PCP) ligand, *wnt11*, is downstream of Nodal signalling ^61^ and *MZwnt11/slb* mutant embryos ^62^ display weaker cell-cell adhesion and a lower blastoderm viscosity than *wildtype* embryos ^18^. Thus, we analysed the dynamics of cell and tissue properties in *MZwnt11/slb* mutant marginal tissues over time and space. Similarly to *MZoep* embryos, the *MZwnt11/slb* marginal tissue exhibits no gradient in connectivity and *α* along the A-V axis (**Fig. 2b, c, Fig. S2a**), which remain close to their corresponding critical points, resulting in the absence of a spatial rigidity pattern (**Fig. 2a, a’, f, Fig. S2b**). Given that *MZwnt11/slb* marginal cells are competent for meso-endoderm specification ^62^, we investigated the effects of the absence of polarized tissue rigidity on Nodal signalling. The fluidized mutants exhibit a higher number of nuclear Smad2-positive cells which are positioned further away from the margin, and the signal is retained longer (**Fig. 2d, e, Fig. S2g, Movie 2**), thereby phenocopying the *Opto-zGrad* embryos. This result supports the hypothesis that Nodal is instructing changes in tissue-scale rigidity by patterning cell-cell adhesion along the A-V axis, mediated via Wnt11 signalling.

If the Nodal/Wnt11-driven rigidity negatively feeds back to terminate Nodal signalling, one would expect to rescue the Nodal signalling dynamics observed in *MZwnt11/slb* mutants, solely by re-establishing tissue rigidity. To test this, we made use of the *Opto-RhoGEF* tool (**Fig. S2e**) ^15,63,64^, to increase cell contractility and reduce the relative surface tension *α* in the most marginal cells, thereby reintroducing a cell-cell adhesion gradient in the *MZwnt11/slb* mutants. Light activation of *Opto-RhoGEF* in the most marginal cells of *MZwnt11/slb* embryos decreases *α* in a graded manner, restoring levels comparable to those in *wildtype* embryos (**Fig. 2c**). This re-introduces a gradient in connectivity (**Fig. 2b, Fig. S2a**) and polarises tissue rigidity towards the YSL (**Fig. 2a, a’, f, Fig. S2b**). Strikingly, the rescue of the tissue rigidity pattern is sufficient to fully rescue nuclear Smad2 spatial and temporal dynamics, resembling those observed in *wildtype* embryos (**Fig. 2d, e, Fig. S2g, Movie 2**).

Altogether, these experiments demonstrate that an adhesion-driven tissue rigidity transition negatively feeds back to Nodal signalling dynamics, in a self-generated manner.

### A rigidity transition restricts Nodal diffusivity

The established regulation of Nodal activity is that Nodal as a short-range activator, induces its own expression and of its antagonist Lefty ^26,32,35,36,47^. Lefty, as a long-range repressor inhibits Nodal activity by diffusing further away and binding to both Nodal and the co-receptors ^26,47,51,65,66^. To understand how tissue rigidity regulates Nodal activity, we explored its potential involvement in the Nodal-Lefty biochemical network (**Fig. 2g**). Theoretical approaches indicate that the geometry of a porous environment through which particles disperse impacts diffusivity ^67–69^. In the context of morphogen transport, the role of tissue architecture has been considered in poro-elastic tissues ^22^, while experimental work proposed that the extracellular fluid structure or porosity can locally change morphogen concentration and diffusivity ^23,27,70^. Furthermore, our recent work showed that tissue rigidification occurs in parallel to drastic changes in tissue porosity ^15^. We therefore hypothesize that the rapid tissue reorganisation occurring during tissue rigidification may directly influence Nodal kinetics within the timeframe of specification. The idea is underlined by the fact that at *α*_*c*_, there is a sudden closure of small interstitial gaps between the cells via the formation of tricellular contacts ^15^ (**Fig. S3a**). This topological change leads to an abrupt collapse of the 3D interstitial fluid network (see Supplementary Theory Note), in which Nodal diffuses, raising the hypothesis that a reduction in tissue porosity may restrict Nodal transport.

Quantification of tricellular contact formation during meso-endoderm specification revealed that in *wildtype* pluripotent tissues with *α* close to *α*_*c*_ most of the potential tricellular contacts are open (**Fig. 3a, a’**). As a result, the interstitial fluid network percolates the 3D-tissue (**Fig. 3a’’**, left panel). However, during specification, the tricellular contacts abruptly close when *α* crosses *α*_*c*_ (**Fig. 3a, a’**), disconnecting the interstitial fluid network (**Fig. 3a’’**, right panel). The relationship between tricellular contact formation and *α* is consistent for all experimental conditions (**Fig. 3a-c, Fig. S3c**), supporting that the topological changes leading to the rigidity phase transition occur under the same conditions at which the geometry of the cell-cell contacts triggers the disconnection of the interstitial fluid network. Quantification of tissue-scale interstitial fluid fraction as a function of *α* revealed that porosity declines even more below *α*_*c*_, after the tricellular contacts are closed (**Fig. 3d**), suggesting that porosity may be further affected by the 3D geometrical changes of multicellular contacts. We mathematically predict that although at *α*_*c*_ the interstitial fluid gaps vanish at tricellular points, they are still present in the central points of tetrahedral cell structures and quadrilateral gaps (**Fig. S3b**, Supplementary Theory Note). By further reducing *α*, we find that at *α* ≈ 0.707 all 3D interstitial fluid gaps close, minimizing tissue porosity (**Fig. 3d**, Supplementary Theory Note). Crucially, when comparing theoretical predictions on tissue porosity against experiments, we conclude that the geometrical changes induced solely by the contact surface tensions can explain the changes in tissue porosity (**Fig. 3d**). The dynamic but drastic changes in tissue architecture occurring during specification suggest that concomitant effects on Nodal transport may arise.

**Figure 3:**
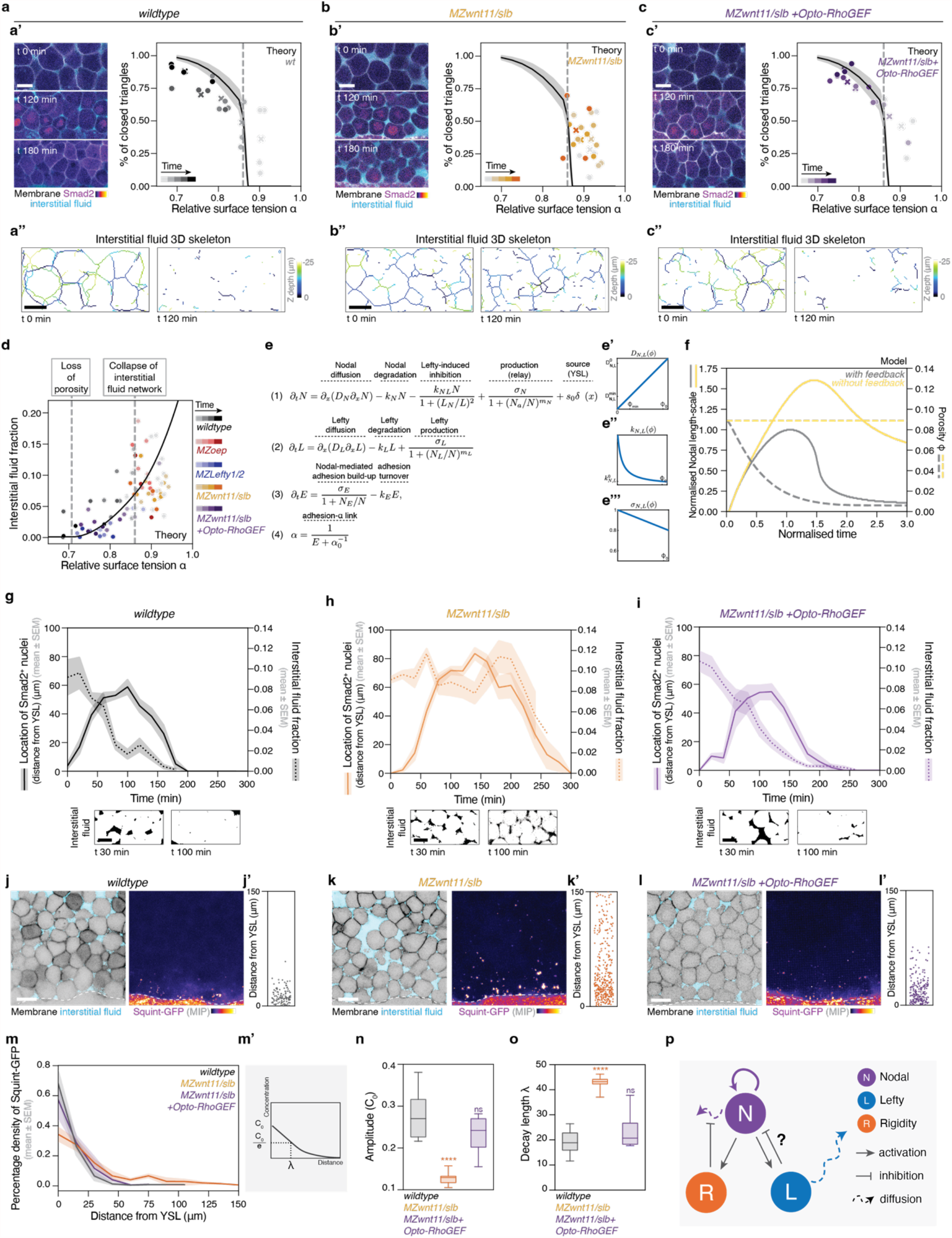
A rigidity-porosity transition negatively feeds back to Nodal signalling by restricting Nodal diffusivity. **(a-c)** (Left) Exemplary 2D confocal sections of the marginal cells close to the YSL over time in *wildtype* **(a’)**, *MZwnt11/slb* **(b’)** and *MZwnt11/slb +Opto-RhoGEF* **(c’)** embryos labelled with membrane-RFP, Smad2-GFP and dextran-647 for interstitial fluid, and (right) quantification of the percentage of closed tricellular contacts as a function of *α* for each experimental condition. Dots indicate single embryos and crosses indicate the mean for each time point. Timepoints are every 30 min. Black curves show simulations of the relationship between *α* and closed tricellular contacts (theoretical results from ^15^). The dotted grey line indicates *α*_*c*_ (n = 4 embryos *wildtype*, n = 4 *MZwnt11/slb*, n = 3 *MZwnt11/slb +Opto-RhoGEF*). **(a’’-c’’)** Exemplary 3D projections of the interstitial fluid skeleton at early and late time points in all experimental conditions, highlighting the changes in interstitial fluid connectivity during the rigidity transition. **(d)** Interstitial fluid fraction as a function of *α*, for all experimental conditions (n = 4 embryos *wildtype*; n = 3 *MZoep*; n = 3 *MZlefty1/2*; n = 4 *MZwnt11/slb*; n = 3 *MZwnt11/slb +Opto-RhoGEF*). Dashed lines indicate the critical points in *α* at which, the tricellular gaps close and the interstitial fluid network collapses (*α*_*c*1_ ≈ 0.866), and all 3D multicellular gaps close and porosity is lost (*α*_*c*2_ ≈ 0.707) (see Supplementary Theory Note). **(e)** Equations implemented to describe the biochemical feedback between Nodal and Lefty, and between local Nodal levels and cell-cell adhesion. Eq. 1 and 2 capture diffusion and degradation of Nodal and Lefty, respectively. Model was adapted from ^71^. Eq. 3 describes the accumulation of adhesion molecules as a response to Nodal levels. Scaling relationships between porosity and diffusion **(e’)**, degradation **(e’’)** and production **(e’’’)** rates (see Supplementary Theory Note). **(f)** Interstitial fluid fraction and Nodal range as a function of time from simulations with (black) and without (yellow) the rigidity feedback. Time is normalised to the time of maximal Nodal length-scale in simulations with feedback. **(g-i)** Quantification of Nodal range (left y-axis, continuous line) and interstitial fluid fraction (right y-axis, dashed line) for *wildtype* (g), *MZwnt11/slb* (h) and *MZwnt11/slb +Opto-RhoGEF* (i) embryos over time. The panels below indicate exemplary interstitial fluid distributions at the onset and peak of Nodal signalling (*wildtype*: n = 6 for Nodal range, n = 3 for porosity; *MZwnt11/slb* n = 4 for Nodal range, n = 3 for porosity; *MZwnt11/slb +Opto-RhoGEF*: n = 3 for Nodal range, n = 3 for porosity). **(j-l)** (left) Exemplary 2D confocal sections of the most marginal cells in *wildtype* (j), *MZwnt11/slb* (k) and *MZwnt11/slb +Opto-RhoGEF* (l) embryos labelled with membrane-RFP and dextran-647 for interstitial fluid, and (right) maximum intensity projections of the same embryos labelled with Squint-GFP. Dashed lines indicate the YSL. (**j’-l’**) Plots of the distribution of Squint-GFP spots as a function of the distance from the YSL for the depicted embryos in (j-l). **(m)** Plot of the normalised Squint-GFP gradient distribution for the experimental conditions shown in (j-l) (n = 5 embryos for each condition). **(m’)** Schematic diagram of the quantification of the decay length *λ* and amplitude *C*_0_. **(n, o)** Boxplots of the fitted amplitude *C*_0_ and *λ* for each condition. **(p)** Illustration of the Nodal-Lefty-Rigidity network, showing the negative feedback from rigidity to Nodal effective diffusivity. MIP, Maximum Intensity Projection; YSL, Yolk Syncytial Layer. Statistics: (n, o) One-way ANOVA and Dunnet’s multiple comparisons test, compared to *wildtype. MZwnt11/slb* p-value < 0.0001 for (n, o); *MZwnt11/slb +Opto-RhoGEF* p-value = 0.51 for (n), p-value > 0.999 for (o). Scale bars: 20 μm

To investigate how the spatial and temporal dynamics of Nodal could be altered by the dynamic changes in tissue porosity, we developed a theoretical framework that integrates biochemical feedback between Nodal and Lefty based on previous reports ^26,47,71^ together with feedback between ligand dispersal and tissue porosity (**Fig. 3e**). We first obtained scaling relationships between tissue porosity and morphogen diffusion, degradation and production from first principles (**Fig. 3e’-e’’’**, Supplementary Theory Note). To model the response of tissue porosity to Nodal levels we assumed that *α* is a function of adhesion ^15,52^ and that Nodal activates adhesion factors. These assumptions were supported by our experimental data showing that tissues undergoing rigidification also display increased levels of α-catenin, which are Nodal/Wnt11 dependent (**Fig. 3e**, equations (3) and (4), **Fig. S3d, d’**). We further employed the derived relationship between tissue porosity and *α* (**Fig 3d**, Supplementary Theory Note). Based on these assumptions, we found that, in the presence of feedback between Nodal and porosity, the coordinated dynamics of Nodal and porosity are consistent with those seen in *wildtype* embryos: Smad2 turns on and reaches its peak, followed by a sharp drop in porosity and then, a timely termination of the Smad2 signal (**Fig. 3f**, grey lines, **g, Fig. S3e**). In the absence of feedback between Nodal and porosity, the model predicts that while porosity remains constant, Nodal signalling reaches further away from the source and takes longer to switch off (**Fig. 3f**, yellow lines). Importantly, this profile matches the relationship between porosity and Nodal dynamics in the *MZwnt11/slb* embryos, which display little decline of the interstitial fluid fraction, show a larger range of Smad2 and a delay in turning off (**Fig. 3h**). Experimentally, the spatiotemporal Nodal dynamics in *MZwnt11/slb* are fully rescuable when restoring tissue rigidity and porosity dynamics via optogenetic increase of cell-cell adhesion strength (**Fig. 3i**). Overall, these results suggest that during the rigidity transition, a concomitant reduction in tissue porosity regulates Nodal signalling dynamics by restricting the Nodal range.

Given that the model assumes that tissue porosity changes Nodal diffusivity, we experimentally assessed the effects of porosity on Nodal dispersal. To test this, we expressed a fluorescently-tagged version of the Nodal ligand Squint in the YSL ^26,66^ (**Fig. S3f**) and quantified its length-scale. In *wildtype* embryos, which rigidify and have reduced porosity, the Nodal ligand gradient is short-range in agreement to previous reports ^26,47,72^ (**Fig. 3j, j’**). However, in *MZwnt11/slb* embryos where the tissue is more porous, the Nodal ligand becomes long-range (**Fig. 3k, k’**). This phenotype could be fully rescued by optogenetically inducing rigidification in the mutant, reverting Squint to a short-range ligand (**Fig. 3l, l’**). A systematic analysis of the profile of Nodal ligand distribution across experimental conditions revealed an exponential-like decay (**Fig. 3m, Fig. S3g-i**). When we quantified the decay length *λ* and amplitude *C*_0_ (**Fig. 3m’**) we could show that *λ* changes as a function of porosity (**Fig. S3j**) and that more porous and fluid-like embryos (*MZwnt11/slb, MZoep*) display increased *λ* and decreased *C*_0_ when compared to less porous and solid-like embryos (*wildtype, MZwnt11/slb +Opto-RhoGEF, MZlefty1/2*) (**Fig. 3n, o, Fig. S3k, l**). Given that an increased *λ* together with decreased *C*_0_ is a signature of increased diffusivity (see Supplementary Theory Note), the observed changes in the Nodal ligand profile suggest that tissue rigidification primarily restricts Nodal effective diffusivity.

### A tissue rigidity transition times Nodal/Lefty dynamics

Although the shift in the Nodal ligand gradient can explain how rigidity restricts the length-scale of Nodal signalling, it is yet unclear how it restricts its time-scale. To this end, we asked whether the porosity-dependent changes in the Nodal length-scale may in turn modify the temporal dynamics by which Nodal transcriptionally activates its downstream targets ^73^, including its inhibitor Lefty (**Fig. 3p**). To investigate this idea, we returned to the theoretical model and we first evaluated the shape of the Nodal signalling gradient, which defines the transcriptional response, with and without feedback. Assuming that both tissues of low and high porosity receive the same initial concentration of Nodal, the model predicts that in the presence of feedback, Nodal activity should form a sharper signalling gradient with higher amplitude at the source (**Fig. 4a**). In contrast, in the absence of feedback, at the same time-scales, Nodal activity should form a shallower signalling gradient with lower amplitude (**Fig. 4b**). To test the predictions of the model, we quantified Nodal activity in *wildtype, MZwnt11/slb* and *MZwnt11/slb* +*Opto-RhoGEF* embryos, by measuring the Smad2 N/C ratio (**Fig. 4c-e**). Accordingly, the cells closer to the YSL in *wildtype* and *MZwnt11/slb +Opto-RhoGEF* embryos displayed a higher peak in Nodal activity, compared to *MZwnt11/slb* embryos (**Fig. 4c-e**, arrowheads, **f**, asterisks). In addition, cells further away from the YSL had higher Nodal activity in the *MZwnt11/slb* embryos (**Fig. 4g**, asterisks), forming a shallower activity gradient (larger *λ*) (**Fig. 4c-e**, dashed line). These findings show that the effects of the rigidity on the ligand gradient directly impact the amount of Nodal cells receive. A logical consequence is that the transcriptional response downstream of Nodal may differ between rigid and floppy tissues.

**Figure 4:**
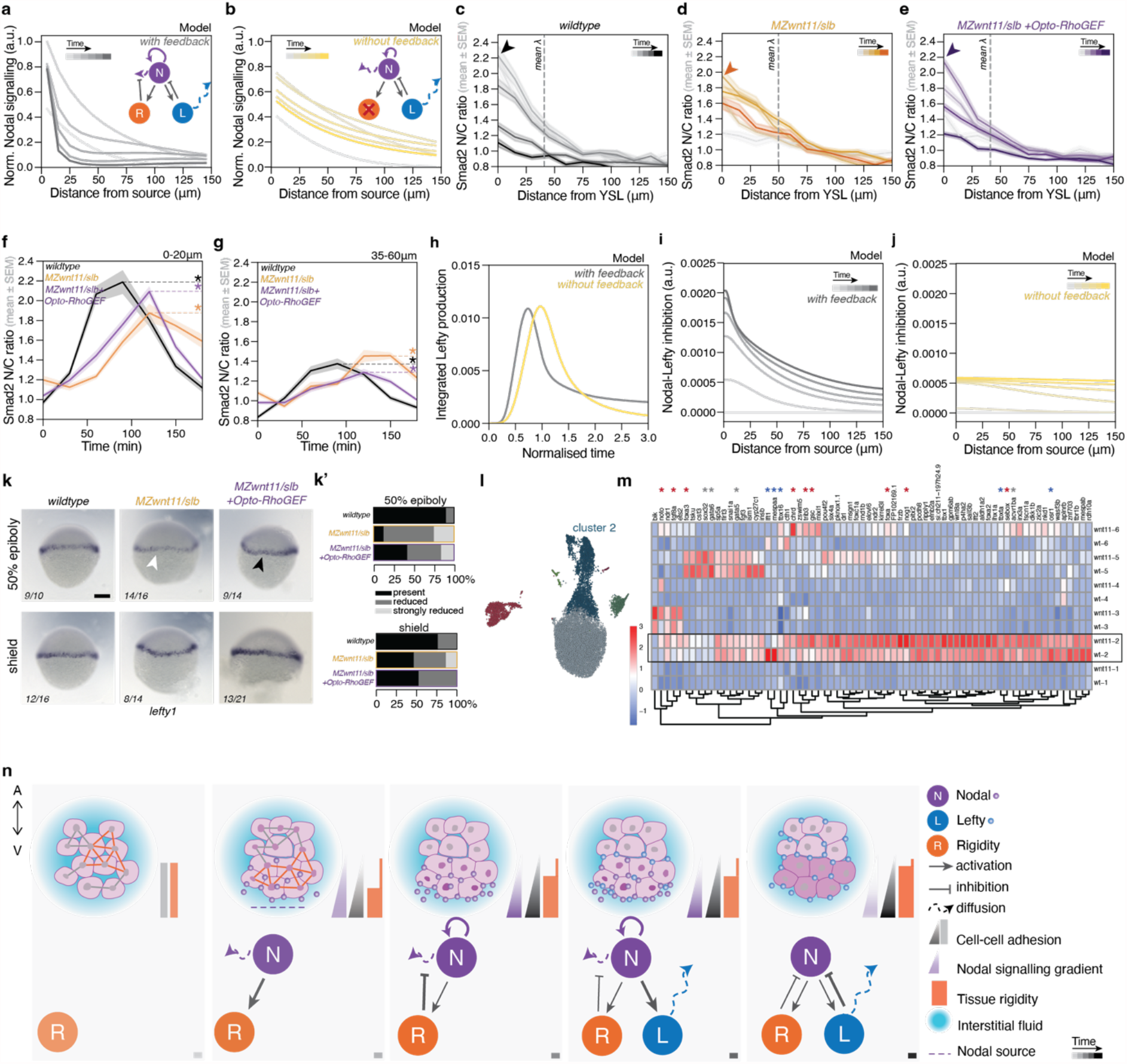
A rigidity transition enhances the Nodal-Lefty biochemical network. **(a, b)** Simulations of Nodal signalling activity in conditions with rigidity feedback (a) and without (b) over space and time. Time points are from 0.25 to 2.75 of normalised time as shown in Fig. 3f **(c-e)** Smad2-GFP N/C ratio signal in *wildtype* (c), *MZwnt11/slb* (d) and *MZwnt11/slb +Opto-RhoGEF* (e) embryos as a function of their distance from the YSL. Timepoints are every 30 min starting from t 90 min. Dotted lines indicate the mean of t 90 min, t 120 min and t 150 min: *wildtype* = 40.8 μm, *MZwnt11/slb* = 49.7 μm, *MZwnt11/slb +Opto-RhoGEF* = 41.2 μm (n = 3 embryos for each condition). Arrowheads indicate the amplitude. **(f, g)** Quantification of Smad2-GFP N/C ratio signal for the conditions shown in (c-e) at the cells close (0-20 μm) (f) and further away from the YSL (35-60 μm) (g) over time. Asterisks indicate Nodal maximum levels. **(h-j)** Simulations of the integrated Lefty production over time (h) and the Nodal-Lefty inhibition over space and time (I, j) with and without the rigidity feedback. Time points are from 0.25 to 2.75 of normalised time as shown in Fig. 3f. **(k)** Representative brightfield images of *lefty1* expression visualized via *in situ* hybridisation in *wildtype, MZwnt11/slb and MZwnt11/slb +Opto-RhoGEF* at 50%-epiboly and shield-stage. The number of embryos with similar phenotype is indicated in the figure. **(k’)** Percentage of embryos showing *lefty1* expression, reduced expression, strongly reduced expression at 50% epiboly-stage and shield-stage. **(l)** UMAP of cells derived from *wildtype* and *MZwnt11/slb* embryos at shield stage profiled with scRNA-seq, coloured by cluster. Clusters were defined in UMAP space, and cluster 2 indicates the meso-endoderm. **(m)** Expression levels for a set of Nodal target genes across all clusters in *wildtype* and *MZwnt11/slb* embryos, with most genotype-specific differences concentrated in meso-endoderm (box, cluster 2). Red asterisks indicate upregulated genes, blue asterisks indicate downregulated genes and grey asterisks indicate unchanged gene expression. **(n)** Schematics illustrating a closed feedback between Nodal signalling and tissue rigidity. N/C, Nuclear-to-cytoplasmic ratio. Scale bar: 200 μm (k)

Given that a crucial downstream target of Nodal is its inhibitor Lefty, which is important for terminating Nodal signalling, we theoretically explored the dynamics of Lefty production when tissue porosity is, or is not, Nodal-dependent. We observe that rigid and less porous tissues which display enhanced Nodal activity near the margin, also display *lefty* production earlier than in fluid and highly porous tissues (**Fig. 4h**). This could in principle accelerate the termination of Nodal signalling, since a source of Nodal degradation is Lefty ^26,35,36,51^. To test this, we quantified the strength of the inhibition of Nodal by Lefty in the model which is predicted to be higher and faster in rigid and less porous tissues than in fluid and highly porous tissues (**Fig. 4i, j**). This prediction suggests that tissue rigidity effectively amplifies and expedites the Nodal-Lefty negative biochemical feedback. To test the model predictions, we performed *in situ* hybridization for *lefty1* RNA in *wildtype, MZwnt11/slb* and *MZwnt11/slb +Opto-RhoGEF* embryos during specification (**Fig. 4k, k’**). This revealed that *lefty1* is not sufficiently expressed early during specification in the fluidized embryos and starts to catch up at later stages (**Fig. 4k, k’**, white arrowhead). The delay in *lefty1* expression is not due to a general developmental delay in the mutant embryos since optogenetic rescue of tissue rigidity partially restores the early expression of the antagonist (**Fig. 4k, k’**, black arrowhead). These results indicate that tissue rigidification regulates the timing of *lefty1* expression, and consequently, the strength of the Nodal-Lefty negative biochemical feedback. In this way, Nodal enhances its own degradation biochemically, by inducing its own inhibitor, and mechanically by reducing its diffusivity and speeding up the induction of its inhibitor.

Self-enhanced morphogen negative regulation was previously shown to endow robustness to morphogen gradients ^74,75^, raising the hypothesis that the above-described double-negative feedback may be important for robust gene expression. To address the downstream consequences of uncoupling the Nodal-Lefty biochemical network dynamics from the tissue rigidity transition, we performed single-cell RNA sequencing in *wildtype* vs *MZwnt11/slb* mutant embryos (**Fig. 4l**). We identified the Nodal-module using two approaches. First, we identified genes whose expression correlated with a curated list of Nodal targets in meso-endoderm cells ^76^ and examined expression levels of genes in this module across cell types (**Fig. 4m**). Second, we performed unsupervised co-expression analysis of all expressed genes in meso-endoderm by reducing dimensionality of transposed cell x gene matrix to cluster genes into modules ^77^. In this gene embedding space, we found a cluster of co-expressed genes that contained all curated Nodal targets, and this module was intact in both *wildtype* and *MZwnt11/slb* embryos (**Fig. S4a**). This data set confirmed that *lefty1* expression is downregulated in the fluidized *MZwnt11/slb* embryos (**Fig. 4m, Fig. S4b**), but also several Nodal targets are upregulated, including, *fgf8a, gsc, chrd* and *noto* (**Fig. 4m**, red asterisks). Notably, several Nodal targets were unaffected (*gata5, sox32*) (**Fig. 4m**, grey asterisks) or even downregulated (*tbxta, tbx16*) (**Fig. 4m**, blue asterisks) suggesting that the meso-endodermal patterning appears fragile in *MZwnt11/slb* embryos, reminiscent of the phenotype of *MZlefty1/2* mutants ^51^. These findings are consistent with the change in the shape of the Nodal signalling gradient in the absence of the rigidity feedback. Shallower gradients with smaller amplitudes imply that cells closer to the source may not receive sufficiently high levels of Nodal to activate genes that are transcribed at high Nodal thresholds (**Fig. 4c-d**, arrows, **f**, asterisks). In contrast, because Nodal decays slower in fluidized embryos, cells further away from the source experience higher levels of Nodal than in the rigid embryos resulting in the over-transcription of genes that require low levels of Nodal (**Fig. 4g**, asterisks). Therefore, the changes in the Nodal signalling gradient induced by the rigidity transition are directly impacting gene expression, scrambling positional information. All together, these results propose the negative feedback from emergent tissue properties as a novel mechanism for timing the Nodal/Lefty reaction-diffusion dynamics to ensure precise patterning of the meso-endodermal domain (**Fig. 4n**).

## DISCUSSION

By uncovering the mechanisms coupling the non-linear dynamics of a tissue rigidity transition to the Nodal/Lefty reaction-diffusion dynamics, this work demonstrates that collective tissue properties shape morphogen signalling gradients. The regulation of the timing of cell signalling dynamics by emergent mechanics unveils a novel mechanism for coordinating a position-appropriate gene expression program to the concomitant tissue shape changes.

Nodal signalling is a paradigm for morphogen-mediated patterning since it displays signatures of wide-spread mechanisms of morphogen gradient formation, including hindered diffusion, the action of signalling feedback and relay, and receptor sensitivity and binding ^26,47,49,65,66,78^. Our finding that the Nodal morphogen gradient is regulated by tissue rigidity provides new insights into how morphogen gradients form and function. For instance, in the case of hindered diffusion, Nodal diffusivity was shown to be slower at the cell-cell interface and faster in cell-fluid interfaces ^70^. We find that porosity drops drastically at the critical point of the rigidity transition, at which cell-fluid interfaces vanish due to the closure of the multicellular contacts. Therefore, this transition may switch Nodal diffusivity from a fast to a slow effective diffusion mode. Given that Nodal at the cell-cell interfaces exhibits higher bound fractions ^70^, it is likely that a sharp drop in porosity locally enhances Nodal signalling activity by facilitating local production and Nodal relay to the immediate neighbours ^47,72,79^ to shape the gradient. Accordingly, our numerical simulations capture the Nodal-Nodal and Nodal-Lefty interactions and the impact of porosity on this biochemical network, when varying the relative contribution to Nodal dispersal via relay (see Supplementary Theory Note). This suggests that porosity can restrict local diffusivity and have similar impact on Nodal dynamics even when transport is mediated primarily via relay. Moreover, although the Nodal activity range is thought to be mostly regulated by the expression of its antagonist Lefty, in the absence of Lefty, Nodal production, although expanded, remains still restricted to a few cell tiers ^51^. A possible explanation for this counterintuitive finding can be drawn from our findings showing that in the *MZlefty1/2* mutants Nodal ligands display low dispersal due to tissue rigidity-mediated restriction (**Fig. S3h**). Thus, the mechanical inhibition of Nodal might act as an early regulator of Nodal diffusion, especially since Lefty translation in zebrafish was proposed to be delayed due to the expression dynamics of miR-430 ^47^. All in all, the effects of the rigidity transition on Nodal diffusivity complements previously proposed mechanisms.

Tissue porosity and the topology of the extracellular fluid compartments, emerge as key regulators of morphogen transport across developing systems. Recent work showed that on the one hand, the topology of the interstitial fluid network can impede morphogen transport, as in the case of Hedgehog proteins ^24^, but on the other hand, facilitate FGF transport ^23^ or even concentrate it in extracellular compartments ^27^. Because biophysical properties of morphogens and of the extracellular space likely define how morphogens are transported (e.g., diffusion-vs cell-based), porosity changes could broadly affect not only the range of morphogen activity, but also additional functionalities like scaling or the interpretation of combined morphogen signalling ^71,80-84^. Moreover, our finding that porosity is non-linearly regulated via material phase transitions can provide simple solutions for limitations of the hindered diffusion-based models. This would be the case when physical boundaries are absent and a morphogen could leak out of the tissue of interest ^85,86^. We show that when the tissue undergoes a rigidity transition, a material boundary effectively appears at the same time. The emergence of such solid-like highly-connected cellular boundary could act as a barrier that impedes morphogen dispersal within the interstitial fluid. Given that the material boundary is also secluding the specifying cells, it may further offer mechanical isolation of the specification zone from neighbouring tissues facilitating germ layer segregation ^87,88^.

Finally, diffusive molecules, like morphogens, may be ideal physiological regulators of control parameters for tissue phase transitions –such as adhesion, cell motility, shape or connectivity– since they can generate graded changes in cell states. Morphogen concentration can induce subtle changes in potential control parameter values between neighbouring cells in an ordered manner in space and time to facilitate patterning refinement ^19,89,90^. Our findings show that when morphogen gradients directly regulate cell control parameters, they trigger symmetry breaking events in space, and rapid transitions in collective tissue organization in time. The closed spatiotemporal feedback between tissue organization and morphogen gradients therefore proposes a new conceptual framework of how patterning and tissue-scale physical properties interact: Morphogen-driven emergent tissue properties act as a design principle that allows morphogens to quickly shape their own length-scales, ensuring that cell fate specification is kept in sync with the rapidly-changing tissue architecture. Last, patterning close to critical points opens a realm of novel mechanisms associated with signatures of critical phenomena ^91^, beyond spontaneous order or symmetry breaking, such as the emergence of bistability in cell mechanical properties after crossing a dynamical bifurcation point ^92^. In this way, some emergent mechanical properties related to dynamical behaviours may be key for understanding phenomena like history-dependent cell decisions, signal sensitivity and adaptation to changing environments ^74,93,94^.

## ACKNOWLEDGEMENTS

We thank Takashi Hiiragi, Caren Norden, Alexandra Schauer, Jaime Agudo-Canalejo, Anna Erzberger, James Briscoe, Caroline Hill and members of the Petridou group for technical advice, critical discussions and feedback on the manuscript; and the Advanced Light Microscopy Facility, Flow Cytometry Core Facility, Gene Core Facility, the Center of Bioimage Analysis and the zebrafish facility at the European Molecular Biology Laboratory (EMBL) for continuous support. We thank the Heisenberg lab for providing fish lines and plasmids. This work was supported by the Weave project “Tissue material phase transitions and their role in embryo pattern formation” from the Deutsche Forschungsgemeinschaft (DFG, German Research Foundation, 518354236, PE 3800/1-1) to N.I.P. and the Österreichischer Wissenschaftsfonds (FWF, Austrian Science Fund, I6533) to B.C-M. N.I.P. was further supported by the European Union (European Research Council Starting Grant 101162743). B.C-M. acknowledges the support of the field of excellence “Complexity of life in basic research and innovation” of the University of Graz. D.K., H.B. and Z.H. were supported by the Francis Crick Institute, which receives its core funding from Cancer Research UK, the UK Medical Research Council, and Wellcome Trust.

## AUTHOR CONTRIBUTIONS

N.I.P. designed the research with substantial input from B.C-M., Z.H., and C.A.. B.C-M. and Z.H. designed the theoretical approaches. C.A. performed all experiments and analysed the experimental data, with help from L.H., C.P-C., V.P., and M.D. on the single-cell transcriptomics. B.C-M. developed the theoretical model linking tissue rigidity and porosity, and E.F. and B.C-M. performed its simulations. D.K., and Z.H., developed the theoretical model of Nodal/Lefty dynamics and its coupling with porosity and D.K., H.B., and Z.H., performed relevant simulations and analysis. N.I.P. wrote the manuscript with substantial input from B.C-M., Z.H., and C.A..

## DECLARATION OF INTERESTS

The authors declare no competing interests.

Exemplary time lapse movies from the margin of *wildtype, MZoep, MZlefty1/2, MZwnt11/slb* and *MZwnt11/slb +OptoRhoGEF* embryos labelled for Smad2-GFP and Dextran-647 for interstitial fluid. Magenta arrowheads indicate Smad2 positive nuclei, white open arrowheads indicate open interstitial gaps (high porosity) and filled white arrowheads indicate close interstitial gaps (low porosity). The dashed line indicated the length-scale of Nodal signalling. Scale bar: 25 μm

## SUPPLEMENTARY FIGURES, FIGURE LEGENDS AND MOVIE LEGENDS

**Figure S1:**
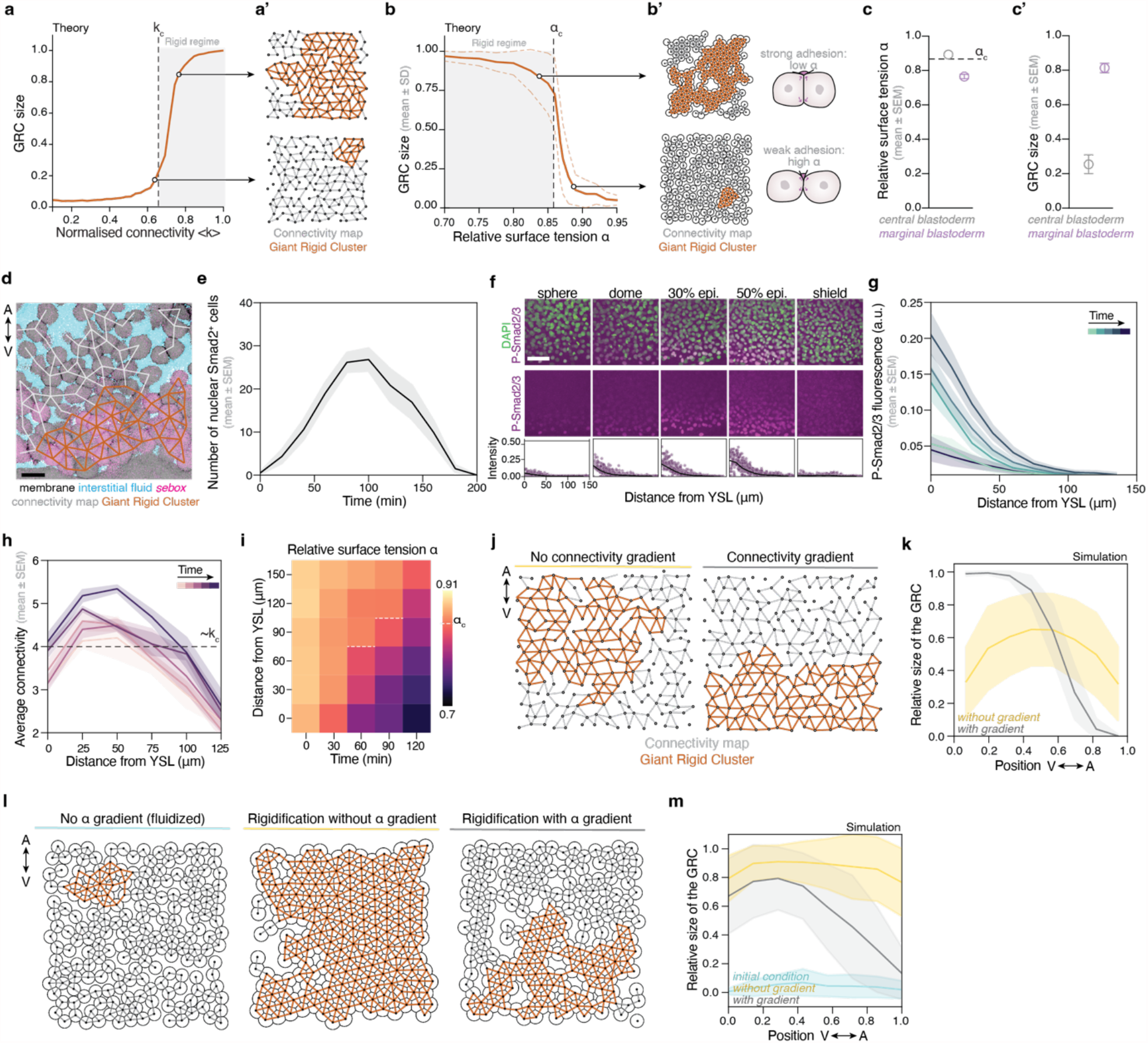
Dynamics of adhesion- and connectivity-dependent tissue rigidity transition during meso-endodermal fate specification in zebrafish. **(a)** Plot of the fraction of the network occupied by the GRC as a function of normalised connectivity <*k*> for simulated random networks and **(a’)** exemplary simulated networks with connectivity above and below the critical point (adapted from ^13^). Rigidity emerges at *k*_*c*_. **(b)** Plot of the fraction of the network occupied by the GRC as a function of the relative surface tension *α* and **(b’)** numerical simulations and overlaid connectivity and rigidity maps of cell arrays with *α* below or above *α*_*c*_ (adapted from ^15^). Rigidity emerges at *α*_*c*_. **(c-c’)** Plots of the relative surface tension *α* (c) and GRC (c’) in the central vs marginal blastoderm at the onset of epiboly (n = 3 embryos each). **(d)** Exemplary confocal section from a transgenic embryo for the meso-endodermal marker *sebox* (Tg(mezzo:eGFP) labelled with membrane-RFP and dextran-647 in the interstitial fluid at t 90 min, with overlaid connectivity and rigidity map, showing the border of the rigid domain coinciding with the border of the specification zone. **(e)** Plot of the number of nuclear Smad2-positive cells as a function of time for the experiments shown in Fig. 1h in a 180 μm wide region. **(f)** Representative maximum intensity projections of fixed embryos stained for DAPI and P-Smad2/3 from blastula to gastrula stages (top) and exemplary quantifications of the P-Smad2/3 signal intensity normalised to DAPI as a function of the distance from YSL of the embryos shown above (bottom). **(g)** Same quantification as in (f) but averaged over several embryos for the reported stages (sphere n = 13, dome n = 9, 30% epiboly n = 14, 50% epiboly n = 16, shield n = 14 embryos). **(h)** Plot of the distribution of average connectivity as a function of the distance from the YSL, over time (n = 4 embryos). Note that connectivity values at the edge of the network (0-25 μm and 125-150 μm) appear lower due to boundary effects (see Methods). The dashed line indicates *k*_*c*_ value for large networks. **(i)** Heat map of the relative surface tension *α* values of *wildtype* embryos over time and space. White dashed line shows the crossing of *α*_*c*_ in space (n = 8 embryos). **(j)** Exemplary simulations showing the location of the GRC without and with a gradient in connectivity along the A-V axis. **(k)** Plot of the distribution of the GRC as a function of the A-V axis for the networks shown in (j). **(l)** Exemplary simulations of cell arrays with overlaid connectivity and rigidity maps. Simulations have been performed starting with connectivity below the critical point and at *α* ≈ 0.95, above *α*_*c*_ (left) and, then, decreasing *α* either without (middle) or with a gradient (right) along the A-V axis. Notice that, for these densities, rigidity is triggered by the decrease of *α*. As seen in the case of hard spheres (blue line), at high *α* regimes the GRC is almost negligible. **(m)** Plot of the distribution of the GRC as a function of the A-V axis for the conditions shown in (l). The time axis in (g) corresponds to the stages in (f) light colour, sphere; dark colour, shield. The time axis in (i) corresponds to intervals of 30 min. A-V, Animal-Vegetal axis; GRC, Giant Rigid Cluster; YSL, Yolk Syncytial Layer. Scale bars: 20 μm (d), 50 μm (f).

**Figure S2:**
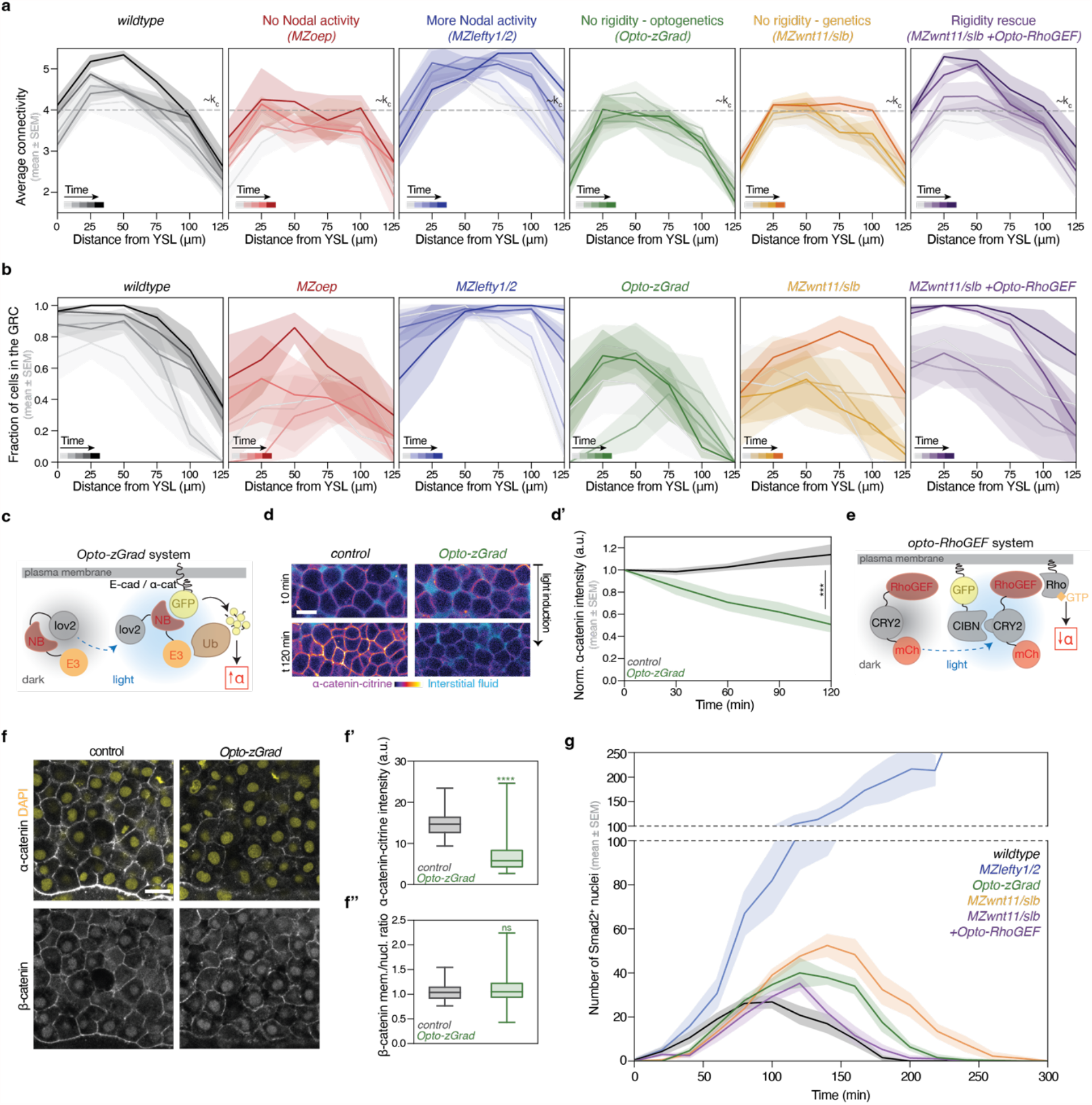
Genetic and optogenetic manipulations of Nodal signalling and tissue rigidity. Plots of average connectivity **(a)** and GRC size **(b)** as a function of the distance from the YSL over time for the experimental conditions shown in Fig. 2a (n = 4 embryos for *wildtype*, n = 3 for *MZoep*, n = 3 for *MZlefty1/2*, n = 3 for *Opto-zGrad*, n = 4 for *MZwnt11/slb*, n = 3 for *MZwnt11/slb +Opto-RhoGEF*). The dashed lines indicate the *k*_*c*_ value for large networks. **(c)** Schematic diagram of the *Opto-zGrad* system, which is composed of an AsLOV2 (LOV2) domain attached to a GFP nanobody (NB) and a domain for proteasomal degradation (E3). Upon light illumination, the GFP-NB is uncaged, so it can bind to GFP and GFP-related proteins (adapted from ^15^). **(d)** Exemplary confocal images of the *Opto-zGrad* tool injected in a zebrafish knock-in line for α-catenin-citrine and **(d’)** quantification of the degradation of endogenous α-catenin upon blue light illumination (n = 5 embryos for control, n = 5 for *Opto-zGrad*). **(e)** Schematic diagram of the *Opto-RhoGEF* tool, which combines a membrane-tagged CIBN and RhoGEF-CRY2. Upon light illumination, CIBN binds to CRY2, translocating RhoGEF to the membrane (adapted from ^15^). **(f)** Exemplary 2D confocal sections of *wildtype* and *Opto-zGrad* injected embryos immunostained for DAPI and β-catenin, and quantifications of **(f’)** α-catenin-citrine at the membrane and **(f’’)** β-catenin membrane-to-nucleus ratio fluorescent intensities (for *wildtype* and *Opto-zGrad*: N = 4 embryos each, n = 80 cells). **(g)** Plot of the number of nuclear Smad2-positive cells as a function of time for the experimental conditions shown in Fig. 2d (n = 6 embryos *wildtype*, n = 3 *MZlefty1/2*, n = 3 *Opto-zGrad*, n = 4 *MZwnt11/slb*, n = 3 *MZwnt11/slb +OptoRhoGEF*). The time axes shown in (b) and (c) correspond to time intervals of 30 min. A-V, Animal-Vegetal axis; GRC, Giant Rigid Cluster; YSL, Yolk Syncytial Layer. Statistics: (d) Unpaired t-test at t 120 min, p-value = 0.0003. (f’) Unpaired t-test, p-value < 0.0001. (f’’) Unpaired t-test, p-value = 0.18. Scale bars: 20 μm (d, f)

**Figure S3:**
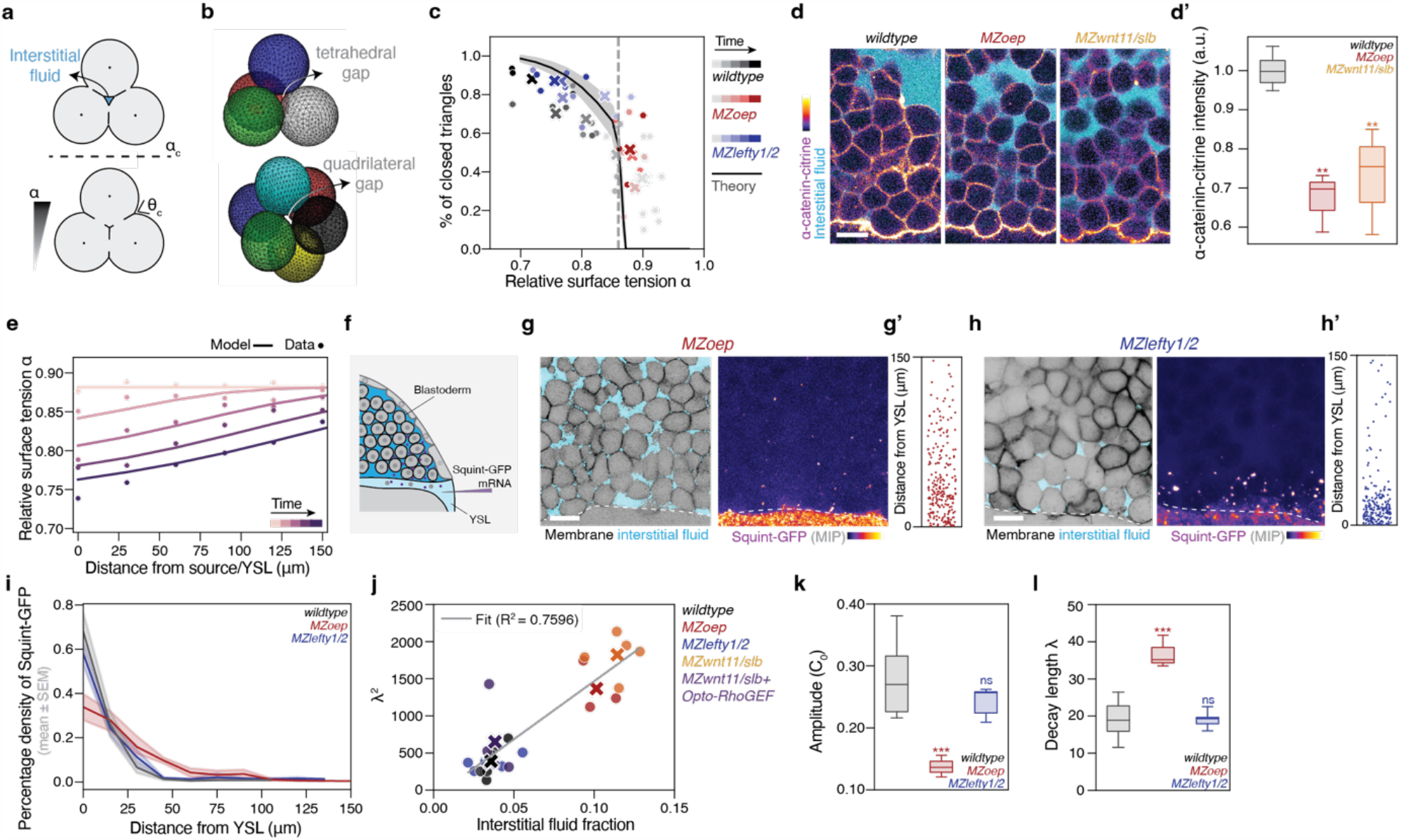
Dependency of Nodal ligand dispersal on tissue porosity. **(a)** Schematic of the closure of the interstitial fluid pocket (cyan) upon crossing of *α*_*c*_ in a 2D cell triplet, with the formation of a tricellular contact and disappearance of the interstitial fluid pocket (adapted from ^15^) **(b)** Schematic of 3D cell topologies showing open tetrahedral (up) and quadrilateral (bottom) gaps. **(c)** Quantification of the percentage of closed tricellular contacts as a function of *α*. The black curve shows the simulated results of the relationship between *α* and the closed tricellular contacts (n = 4 embryos for *wildtype*, n = 3 for *MZoep*, n = 3 for *MZlefty1/2*). **(d, d’)** Representative images of the endogenous α-catenin-citrine signal at t 120 min in *wildtype, MZoep* and *MZwnt11/slb* embryos at the margin, labelled for interstitial fluid by Dextran-alexa647 and quantifications of the endogenous α-catenin-citrine signal, all normalised to the *wildtype* (n = 7 embryos *wildtype*; n = 3 *MZoep*; n = 7 *MZwnt11/slb*). **(e)** Spatial profiles of *α* from simulations with feedback, overlaid with experimental values from Fig. 2c (*wildtype*). **(f)** Schematic of the YSL injection of Squint-GFP mRNA for ligand diffusion assay. **(g, h)** Exemplary 2D confocal sections of the most marginal cells in *MZoep* (g) and *MZlefty1/2* (h) embryos labelled with membrane-RFP and dextran-647 for interstitial fluid, and (right) maximum intensity projections of the same embryos labelled with squint-GFP. Dashed lines indicate the YSL. **(g’-h’)** Plots of the distribution of the Squint-GFP spots as a function of the distance from the YSL for the depicted embryos in (g-h). **(i)** Plot of the normalised Squint-GFP gradient distribution for *wildtype, MZoep and MZlefty1/2*. (n = 5 embryos *wildtype*; n = 4 *MZoep*; n = 5 *MZlefty1/2*). **(j)** Plot of the decay length *λ* as a function of their respective interstitial fluid fraction. The black line shows a linear fit with R^2^ indicated in the plot (numbers same as Fig. 3m and S3i) **(k-l)** Boxplots of the fitted amplitude *C*_0_ and decay length *λ* for each condition (numbers as in (i)). For (c) and (j), dots indicate single embryos and crosses indicate the mean for each condition. Timepoints are every 30 min. Abbreviations: MIP, Maximum Intensity Projection; YSL, Yolk Syncytial Layer. Statistics: (e, k, l) ANOVA and Kruskal-Wallis multiple comparisons test, compared to *wildtype*: *MZoep* p-value = 0.007, *MZwnt11/slb* p-value = 0.008 for (e); *MZoep* p-value = 0.001, *MZlefty1/2* p-value = 0.47 for (k); *MZoep* p-value = 0.0008, *MZlefty1/2* p-value > 0.99 for (l). Scale bars: 20 μm

**Figure S4.**
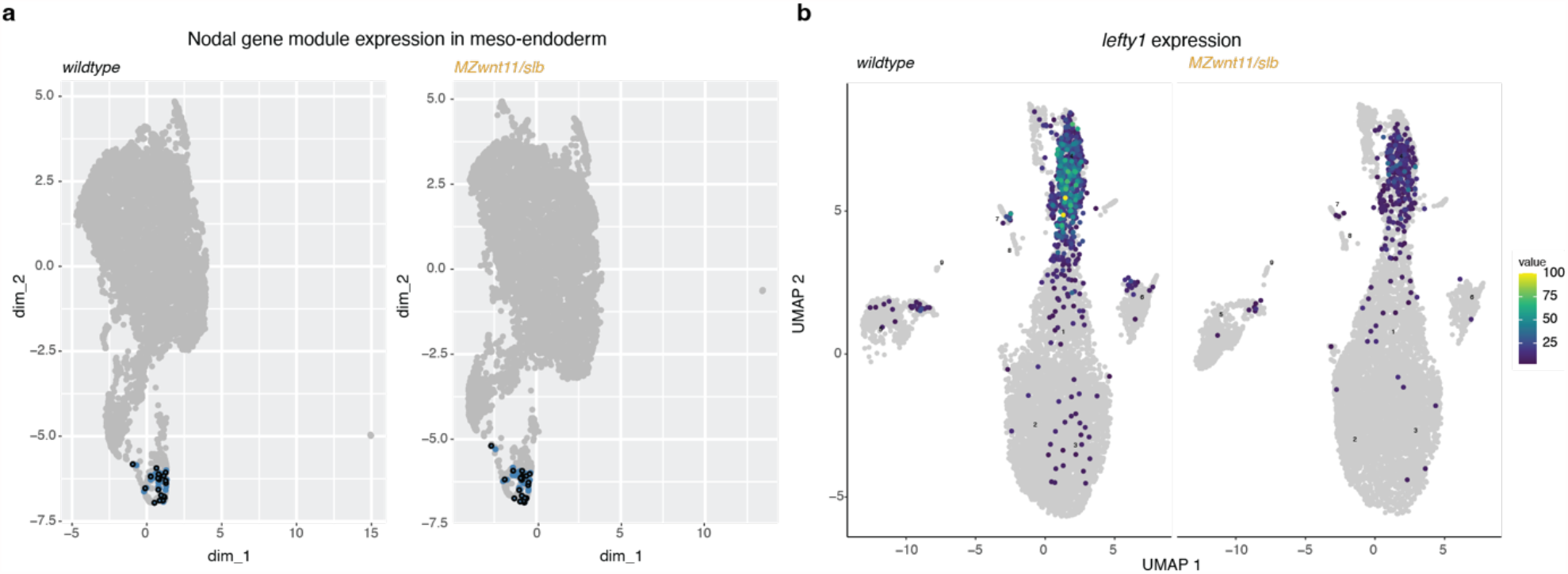
Gene expression analysis with and without the rigidity transition. **(a)** Gene module UMAP, where each point represents a single gene and genes are grouped based on their co-expression in meso-endoderm cells. Known Nodal target genes (highlighted in blue) co-localize in the gene module UMAP, reflecting a shared co-expression pattern that is intact in both *wildtype* (left) and *MZwnt11/slb* (right) shield stage embryos. **(b)** *lefty1* expression in *wildtype* and *MZwnt11/slb* embryos.

**Supplementary Movie 1. Mapping tissue rigidity during meso-endoderm specification**

Exemplary time lapse movie of a transgenic embryo expressing eGFP under the *sebox* promoter, labelled with membrane, nuclei and interstitial fluid markers, with overlaid rigidity map in a 150-180 μm region including parts of the specification zone and the overlaying pluripotent blastoderm. The black network is the connectivity map and the red network is the Giant Rigid Cluster marking tissue rigidity. Scale bar: 50 μm

**Supplementary Movie 2. Nodal signalling dynamics in different experimental conditions**

## METHODS

### Zebrafish handling

Zebrafish (*Danio rerio*) were raised at 28.5°C under a 14 h light / 10 h dark cycle ^95^. The following zebrafish strains were used in this study: *wildtype* A2B2, Tg(mezzo:eGFP) ^96^, *MZwnt11/slb*^tx226 62^, *MZoep*^tz257 97^, *MZlefty1*^a145^*lefty2*^a146 51^, *Gt(ctnna-citrine)*^*ct3a 98*^. From the above lines, this study generated the following lines: *MZWnt11/slb*^tx226^;Tg(mezzo:eGFP), *MZWnt11/slb*^tx226^;*Gt(ctnna-citrine)*^ct3a^, *MZoep*^tz257^;*Gt(ctnna-citrine)*^ct3a^. For raising, *MZoep*^tz257^ embryos were rescued with injection of 200pg of *oep* mRNA and *MZlefty1/2* mutants were rescued by growing the embryos in 2.5 µM of Nodal inhibitor in E3 from 1-cell stage for 24 h (S4696 Sigma-Aldrich, SB505124). Zebrafish embryos were grown at 28.5°C and maintained in 1x Danieau’s medium for all experimental incubations. Staging was performed as previously described ^99^. All animal experiments were carried out according to the guidelines of the Committee for Animal Welfare and Institutional Animal Care and Use (IACUC) under EMBL’s Policy on the Protection and Welfare of Animals Used for Scientific purposes (IACUC code 21-010_HD_NP).

### Embryo microinjections

Zebrafish embryos were injected using glass capillary needles (30-0020, Harvard Apparatus, MA, USA) that were pulled with a P-97 needle puller (Sutter Instrument) and attached to a PV820 microinjector system (World Precision Instruments). mRNA *in vitro* transcription was performed with the mMessage mMachine SP6 kit (Thermo Fisher Scientific, AM1340). The following amounts were injected at the 1-cell stage in the yolk: 70 pg *membrane-RFP* ^100^, 20 pg *histone-BFP* ^101^, 50 pg *Smad2-GFP* ^102^, 35 pg *Opto-Zgrad* ^15^, 70 pg *CIBN-CAAX* ^15^, 15 pg *Opto-RhoGEF2-CRY2* ^15^. Interstitial fluid was labelled with 0.5 nl of 0.6 μg/μl 10000 MW Dextran-Alexa Fluor^®^ 647 (Thermo Fisher Scientific, D22914) injection in the blastoderm at 1K-oblong stages (3-3.5hpf). Ligand YSL injections were performed at sphere stage by co-injecting in the YSL 30pg of *histone-BFP* mRNA, 170 pg of *squint-GFP* mRNA ^26^ and 0.68% phenol red (Sigma, P0290). Phenol red was used to evaluate the success of the injection.

### *In situ* hybridization

*In situ* hybridizations were performed as previously described ^103^. Briefly, 50% epiboly-stage and shield-stage embryos were fixed with 4% paraformaldehyde (PFA) in PBS overnight at 4 ^o^C, then washed in PBS, dechorionated, dehydrated and stored in 100% methanol at −20 °C. Embryos were then rehydrated in PBS, permeabilized in PBT (PBS + Tween20 0.1%) and incubated in hybridization buffer (50% Formamide, 5X SSC, 0.1% Tween20, 50 μg/ml Heparin, 500 μg/ml tRNA, and citric acid) with digoxigenin-labelled RNA probe for *lefty1* overnight (Heisenberg lab). Embryos were washed in serial dilutions of SSC, then incubated overnight with an Alkaline phosphatase anti-digoxigenin antibody (11093274910, Roche), washed and stained with NBT (11383213001, Roche) and BCIP (11383221001, Roche) and the reaction was stopped with 100% ethanol.

### Immunostaining

Embryos were fixed in 4% PFA solution in PBS overnight at 4 ^o^C. The next day embryos were washed 3x 5 min in PBS and manually dechorionated. Samples were washed 5x 5 min in PBSTr (PBS + 1% TritonX-100), blocked 2x 2 h in blocking solution (PBSTr + 10% goat serum + 1% DMSO), and incubated overnight at 4 °C with primary antibody anti-P-Smad2/3 (8828S, Cell Signalling) 1:1000 in blocking solution, as described in ^48^. Samples were washed 5x 10 min in PBSTr, 3x in PBS + 0.1% TritonX-100 (PBTr) for 1 h, and incubated overnight at 4 °C with secondary antibody (1:500). Samples were then washed 2x 5 min in PBTr, labelled with DAPI (1:1000, D1306, Invitrogen) for 20 min and washed 5x 10 min in PBSTr. Embryos were transferred to 80% glycerol in PBS with 0.4% N-propyl-gallate, manually de-yolked with forceps and a tungsten needle tip, flat-mounted on glass slides and covered with a glass cover-slip. β-catenin immunostainings were performed as described above with the difference that they were blocked in blocking solution (PBS + 1% Triton + 2% BSA for 3 h), and anti-β-catenin primary antibody incubation (C7207, Sigma-Aldrich) was performed overnight at 4 °C (1:100 in PBS+0.5% Triton + 2%BSA). The next day the samples were washed 5x 10 min with PBSTr (PBS + 0.5% Triton). Secondary antibody (anti-mouse Alexa-647, 2272554, Invitrogen) incubation was performed at room temperature for 4 h (1:1000 in PBS + 0.5% Triton + 2% BSA).

### Image acquisition

Dechorionated embryos were embedded in 0.6% low melting point agarose (Invitrogen, Cat. No. 16,520-050) on a customized agarose mold in a 60 mm dish (Greiner, 628102). Mounted embryos were kept in an incubation chamber at 28.5°C throughout acquisition. All imaging was performed with an upright confocal microscope Zeiss LSM 980 equipped with Airyscan 2 with Axio Examiner, with a 20x immersion objective W Plan-Apochromat 20x/1.0 Corr DIC M27 75mm. Timelapse z-stack images were acquired with 10-15 min intervals. Images were acquired in ZEN3.3 (blue edition) software (Carl Zeiss).

### Optogenetics

Embryos were kept in the dark until the selected developmental stage. To prevent photoactivation, all sample handling and mounting was performed under red light filters (Lee Filter 106, Primary Red) to block bright-field illumination. For α-catenin-citrine degradation, *Gt(ctnna-citrine)*^*ct3a*^ homozygous embryos were microinjected with opto-zGrad as described above. Opto-zGrad photoactivation and imaging of α-catenin-citrine degradation were carried out with 488 nm light pulses (laser power set between 3-6%, corresponding to an out-of-objective power of 50–104 µW) every ~2.5 min, with a z-stack size of ~100 µm (3.5 µm spacing) for about 30 min, then normal imaging was performed. Dark effects were sometimes observed in a concentration-dependent manner. For Opto-RhoGEF experiments, *wildtype* embryos were microinjected with Opto-RhoGEF-CRY2 and CIBN-CAAX mRNAs and were photoactivated/acquired with a 5-6% 488 nm laser (corresponding to 86-104 µW) in a total z stack of 80 µm (3 µm spacing), with pulses every ~2.5 min, from sphere-onset of doming stage for about 30 minutes, then normal imaging was performed.

### Data analysis and quantification

Acquired data were processed using Fiji ^104^, Imaris (v 10.1, Oxford instruments) and Cellpose v2 and v3 ^105,106^. All the data was analysed and plotted in Python (v 3.10.9) with standard packages: `matplotlib` (v 3.7.0), `pandas` (v 1.5.3), `seaborn` (v 0.12.2), `numpy` (v 1.23.5).

### Sebox fluorescence

Intensity profiles were generated with the `plot_profile()`function in Fiji with lines of thickness 110 μm in width on a maximum intensity projection over a length of 150 μm and binned every 1 μm. The signal for each embryo was then normalised to the maximum across the t 0, t 30, t 60, t 90, t 120 min timepoints from the last YSL division.

### *α*-catenin intensity

The intensity of the endogenous α-catenin fluorescence was quantified in homozygous Gt(ctnna-citrine)^ct3a^ embryos, either in *wildtype* (with and without opto-zGrad), or mutant backgrounds (*MZoep, MZwnt11/slb*). For the optogenetic experiments, α-catenin-citrine intensity was measured in a 150 x 80 µm ROI in the margin of the blastoderm at t 0, t 30, t 60, t 90 and t 120 min and normalised to the t 0 min intensity of each embryo. To compare α-catenin levels between the different genetic backgrounds, fluorescence intensity was measured in a 130 x 65 µm ROI in the margin of the blastoderm at t 120 min and each condition was normalised to the mean of the controls of the same experiment.

### Image segmentation

Blastoderm images were segmented using Cellpose v2 and v3 ^105^, with manual corrections being performed where necessary.

### Reconstruction of connectivity maps and rigidity analysis

Connectivity networks were reconstructed in Fiji on ROIs of 150 x 150 µm cropped from time lapse movies, aligning the margin with the YSL at the bottom of the ROI, every 30 min from the last YSL division. As previously described in ^13,15^, networks were reconstructed on the 2^nd^-3^rd^ layer of deep cells in the tissue. The interstitial fluid channel was binarized in Fiji via thresholding. To obtain more accurate segmentation results, the membrane channel was processed as follows: the “AND” operator in the `Image Calculator` function was used between the membrane channel and the inverted binarized interstitial fluid channel (pixel value 0 for interstitial fluid), to convert the membrane channel areas occupied by interstitial fluid to pixels values of zero. A custom Fiji plugin based on the `region adjacency graph` function from the MorphoLibJ plugin ^107^ was used to reconstruct connectivity networks from the cell segmentations and overlay them to the tissue and segmentation, where label adjacency was evaluated via the presence or absence of neighbouring pixels. Coordinates of the nodes are exported, as well as the adjacency matrix. The reconstruction of the networks and rigidity analysis was performed with a python version of the pebble game algorithm (`pebble.py`) and plotted using matplotlib ^13,15^. Average connectivity was calculated as the total number of contacts (defined as described in connectivity map reconstruction) divided by the total number of cells in the image. Normalised connectivity (*<k>*) was calculated in each confocal section as connectivity divided by the maximum potential connectivity (*k*_*max*_) (computed as described in ^13^). For the rigidity analysis along the A-V axis, the following properties of each node were extracted: *xy* coordinates, number of contacts, and if the node is in the GRC, and were then plotted in 25 µm bins. Average connectivity in this case was calculated as the total number of contacts divided by the number of cells within each bin. Due to cropping the image to perform the analysis, connectivity at the edges of the network is underestimated since some of the neighbours of the cells at the boundaries are out of the field of view. This is relevant for connectivity of the cells at the edges of the network (0-25 µm and 125-150 µm). Please note that the critical point in connectivity for a rigidity transition is ~4 contacts per cell for average connectivity, and ~2/3 of maximum potential connectivity (*k*_*max*_ ≈ 0.667) for normalised connectivity ^13^.

### Relative surface tension *α*

For mapping the spatiotemporal patterns of the contact surface tensions, cell-cell contact angles were measured every 30 min from the last YSL division. Only one angle was measured for each pair of cells, and this was done in the middle of the cell volumes after inspecting the different Z planes. A custom Fiji script was used to extract the *xy* coordinates relative to the YSL, which was marked with the line tool. A custom python code was used to calculate the *α* parameter from the angles in radians with the formula 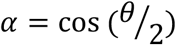, bin the data along the A-V axis every 30 µm, and plot the measurements across space and time. Heatmaps were done with the `seaborn` library based on matplotlib after transposing the data into pivot tables.

### Interstitial fluid fraction

The interstitial fluid fraction was calculated by binarizing the interstitial fluid channel in Fiji the interstitial fluid channel labelled and calculating the fluid fraction in an ROI of 120 x 80 µm.

### Tricellular contacts (closed triangles)

A custom-made Python pipeline for morpho-feature quantification in 2D ROIs of 120 x 70 µm was used ^15^. In brief, segmentation masks were generated with Cellpose and manually curated after visual inspection of the membrane and interstitial fluid pockets. Using `scipy.ndimage.generic_filter` a junction map was generated and the locations where three cells meet were identified. These were then normalised to the number of triangles obtained from the connectivity maps.

### Skeletonization

Skeletonizations of 3D Z-stacks of 25 µm depth of a 100 x 50 µm ROI were performed in Fiji on the interstitial fluid channel after binarization with the `Skeletonize 2D/3D()`function. A maximum projection of the skeleton was done and color-coded by depth.

### Nodal signalling activity

To dynamically measure Nodal activity during development, Smad2-GFP mRNA-injected embryos were imaged via confocal microscopy from sphere-stage for at least 3 h every 10-15 min. A region of 180 µm width was cropped and Smad2 positive nuclei were marked with the spot function in Imaris (Imaris 10.1, Oxford Instruments) at every time point and the *x*,*y* coordinates were extracted. The data was analysed in python with custom scripts: the nuclei *x*,*y* coordinates were normalised to the lowest nucleus along the y-axis. Data points were then binned every 20 min, and aligned at the last YSL division as t 0 min. Length-scale was measured as the maximum distance from the YSL for each timepoint, and the count as the number of positive nuclei for each timepoint. The nuclear-to-cytoplasmic ratio was obtained by measuring in Fiji the mean grey value for the Smad2-GFP channel respectively in the nucleus and in the cytoplasm for each timepoint for a region of 120 µm in width, along the AV axis. The data was analysed in python with custom scripts: the nuclei *x*,*y* coordinates were normalised to nucleus closest to the YSL along the y-axis, and the ratio between nuclear and cytoplasmic signal was performed. The datapoints were then binned every 15 µm and every 30 min, and aligned at the last YSL division as t 0 min. The *C*_*max*_ and *λ* values were obtained by fitting the datapoints with the `curve_fit()` function within the `scipy.optimize` package as 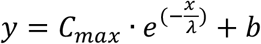, and only fits with R^2^ > 0.75 were taken into account.

### Ligand gradient

The spots detection function in Imaris was used to detect the squint-GFP bright spots from 3D Z-stacks at a timepoint ~ 2 h after injection at sphere stage. The *xy* coordinates of the spots were then extracted and the data was analysed and plotted with custom Python scripts. To obtain the profiles of diffusion, the data was binned along the y-axis every 15 µm, expressed as distance from the YSL, and plotted as density functions, after normalising every bin to the total per embryo, to correct for injection differences. The *C*_*max*_ and *λ* values for each embryo were obtained by fitting the datapoints with the `curve_fit()` function within the `scipy.optimize` package as an exponential decay, with the following function: 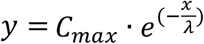. The relative differences in *C*_*max*_ and *λ* values between the experimental conditions were preserved even when quantifying without the normalisation step.

### Single-cell RNA sequencing

ACME (ACetic-MEthanol), a dissociation protocol for single-cell transcriptomics, which simultaneously fixes the tissue, was used to avoid potential effects of the dissociated tissue architecture to gene expression ^108^. Briefly, shield-stage *wildtype* and *MZwnt11/slb* embryos were dechorionated and treated with the ACME solution. Dissociated cells were cryopreserved by adding 10% DMSO and stored at −80 ^o^C. To enrich for singlets and eliminate cell debris and aggregates, ACME-dissociated cells were stained with DAPI and cell sorting was performed by the FACS Facility at EMBL. Sample quality was assessed by evaluating cell morphology using a ZEISS LSM 980 with Airyscan 2 microscope and RNA integrity was verified using a Bioanalyzer. Single-cell RNA sequencing was carried out using the 10x Genomics platform. cDNA libraries targeting ~10,000 cells per sample were prepared using the Chromium Next GEM Single Cell 3′ Reagent Kits v3.1 (10x Genomics) following the manufacturer’s instructions. Sequencing was performed on a NextSeq2000 P2 platform with 50 bp paired-end reads by the Gene Core Facility at EMBL. Reads were processed using CellRanger. Output cell metadata and count matrix were read in and pre-processed in Monocle3 using a standard workflow: `estimate_size_factors()` *->* `preprocess_cds()` with 30 principal components, followed by sample-wise batch correction in PCA space `align_cds(alignment_group = ‘Sample’, residual_model_formula_str = “~log10(n.umi)”)`, dimensionality reduction `reduce_dimension(preprocess_method = ‘Aligned’, max_components` `= 2, reduction_method = ‘UMAP’)`. Cell clustering was performed in the batch-corrected PCA space `cluster_cells(reduction_method = ‘Aligned’)`. Expression signatures for Nodal target genes were generated using `aggregate_gene_expression()` with log normalised of mean expression values across cells. *De novo* identification of gene modules was performed using `find_gene_modules()` on the subset of meso-endodermal cells, separately for *wildtype* and *MZwnt11/slb* mutant samples. Sequencing data have been deposited to NCBI GEO GSE299074, GSM9032491, GSM9032492.

### Statistics and reproducibility

The statistical analyses were performed with GraphPad Prism 10.0. Statistical details of experiments are reported in the figures and figure legends. Sample sizes are provided in the figure legends, and no statistical test was used to determine sample size. The biological replicate is defined as the number of embryos. No inclusion or exclusion criteria, randomization, or blind allocations were applied, and all analyzed samples were included. Unless differently stated in the figure legends, the graphs show mean ± SEM, and the error bars are calculated and shown based on the number of cells or embryos, as indicated. The statistical test used to assess significance is stated in the figure legends and was chosen after testing each group for normality using the Shapiro-Wilk test and for homogeneity of variances using Levene’s test. For comparisons between two groups, a two-tailed Student’s t-test was used for parametric distributions with equal variances. For multiple pairwise comparisons, an ANOVA followed by Dunnett’s test was used for parametric distributions, while a Kruskal-Wallis test followed by Dunn’s test with Dunnett’s adjustment (for pairwise comparisons) was applied for non-parametric distributions.

### Reagents, data and code availability

All reagents, codes and data generated in this study will be publicly available upon manuscript publication.

## Supplementary Note

## 1 Theoretical framework

In this section we lay out the theoretical framework we have developed to couple Nodal-Lefty reaction diffusion dynamics with tissue porosity. We further present derivations that couple the relative surface tension between cells to tissue porosity.

### 1.1 Coarse-grained concentrations in a porous environment

We base our analysis on a model for Lefty and Nodal dynamics in the zebrafish blastula that has been proposed as an inhibitor-activator system and previously fitted to experimental data in [1]. We couple this model with a spatially varying tissue porosity as described below. We model the blastoderm tissue as a continuous 1D domain along the direction of the morphogen gradient, *x ∈* [0, *l*], with *x* = 0 denoting the border with the YSL and *l* the length of the measurement window. We assume that all quantities are averages over the thickness of the tissue layer (3-5 cells thick) and ignore any dependence on the curvature of the embryo, as this remains constant during the time window and in the region of interest.

We introduce coarse-grained concentrations *c*, locally averaged over the extracellular and cellular regions (see Fig SN1). The time evolution of a coarse-grained concentration field *c*(**x**, *t*) is generally given by an advection-diffusion equation [2]

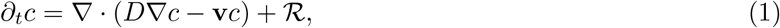

with a diffusive flux −*D ∇ c*, an advection velocity **v**, and reaction terms denoted by ℛ. Although we do not consider hydrodynamic flows, the spatially heterogeneous porosity *ϕ*(**x**) requires a non-zero apparent advection velocity as derived below.

The coarse-grained concentration is defined through the porosity as

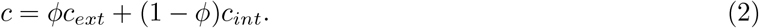

If the extracellular and intracellular concentrations, given by *c*_*ext*_ and *C*_*int*_ respectively, are related by linear reactions (such as endo- and exocytosis with constant rates) and those equilibrate on time scales faster than the tissue-wide morphogen dynamics, we have *c*_*int*_ = *βc*_*ext*_ and

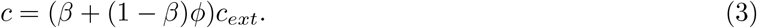

In the absence of reactions at the coarse-grained level, ℛ = 0, at steady state extracellular (and intracellular) concentration must be uniform in space 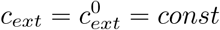. and fluxes must vanish,

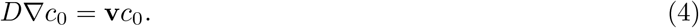

With 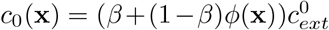, this determines the form of an apparent velocity, resulting from gradients in the porosity *ϕ*(**x**),

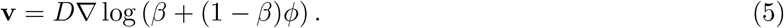

This contribution to the transport equation constitutes an apparent flux towards regions of higher porosity. For *β* = 0, we have **v** = *D∇* log(*ϕ*), which is the result for a porous medium with impermeable obstacles [3]. Conversely, for *β* = 1, i.e. *c*_*int*_ = *c*_*ext*_, the velocity term vanishes.

### 1.2 Coupled dynamics of Lefty and Nodal

Starting from equations (1) and (5) for the concentrations of Nodal *N*(*x*) and Lefty *L*(*x*) we now add the reaction terms following the inhibitor-activator interactions [1]. We then arrive at a coupled system of partial differential equations,

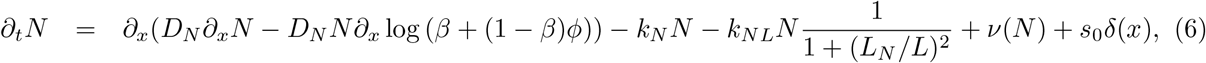

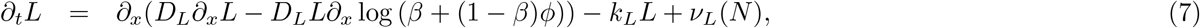

**Figure SN1:**
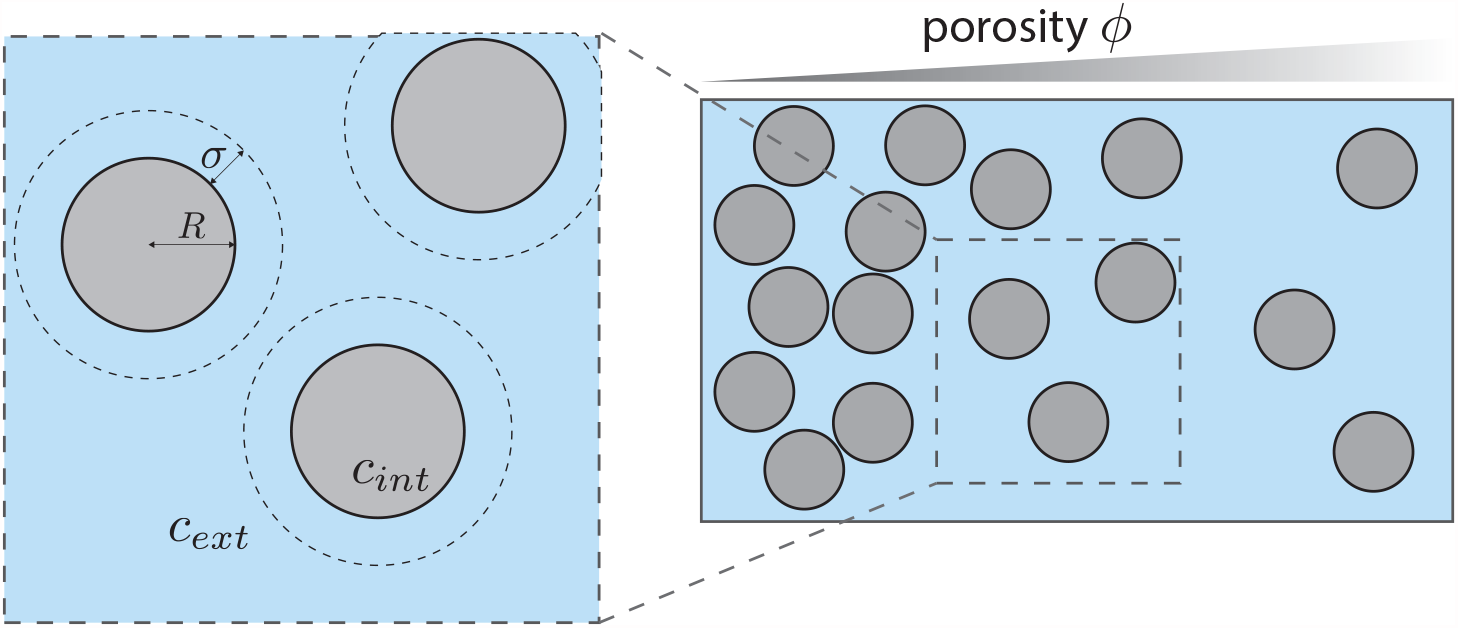
We model the tissue as a porous material with spatially graded porosity *ϕ*(**x**). Cells are denoted by grey circles and assumed to have a constant radius *R*. The porosity is the locally-averaged fraction of extracellular space, e.g. in an small region of the tissue having area *A*_*c*_ containing *N*_*c*_ cells it is given by Equation (11). We further denote by *c*_*ext*_ and *c*_*int*_ the extracellular and intracellular concentrations of morphogen, which are locally averaged to obtain *c* (see Eq. (2)). The morphogen uptake into a cell happens from a ring of size *σ* around the cell.

where both morphogens diffuse with diffusivities *D*_*N*,*L*_(*ϕ*) and are degraded linearly with rates *k*_*N*,*L*_(*ϕ*) [4]. We also include Lefty-mediated inhibition of Nodal (*k*_*NL*_), self-activation of Nodal via relay [5] (ν(*N*)) and Nodal-mediated Lefty activation (ν_*L*_(*N*)) [6, 7]. There is an actively regulated influx of Nodal molecules from the YSL into the tissue which we model by a delta-peak of Nodal production at *x* = 0 with a magnitude *s*_0_. At the tissue boundaries the diffusive flux is vanishing,

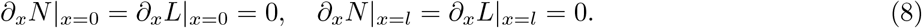

For the activation terms we assume a Hill-type function

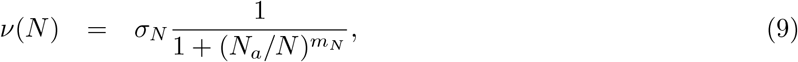

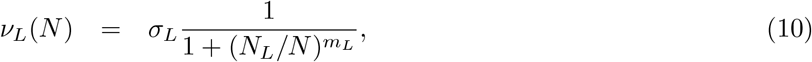

with activation strengths *σ*_*N*,*L*_, Hill coefficients *m*_*N*,*L*_ and activation thresholds *N*_*a*,*L*_. We assume that the coarse-grained concentrations used here capture signalling levels, e.g. as measured by Smad2-positive nuclei (see main text Fig. 2 d,e).

### 1.3 Scaling of degradation rates and diffusion constants with tissue porosity

Here we consider a 2D projection of the tissue. The porosity *ϕ* = *ϕ*(**x**) is defined as

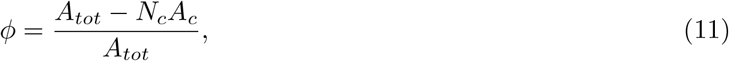

that is the fraction of extracellular space in a small region of the tissue of area *A*_*tot*_ centred at **x** and containing *N*_*c*_ cells of area *A*_*c*_. Note that this 2D porosity is equivalent to the interstitial fluid fraction (IFF) measured in the experiments throughout the article. From the literature on porous materials we expect that porosity affects the coarse-grained transport properties of morphogens in a tissue [8, 9, 10].

For many types of porous materials it has been shown that effective diffusivity follows an approximately linear scaling with porosity

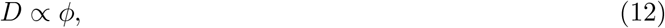

for large and intermediate porosities [8, 3, 11]. Deviations from this scaling are possible at small porosities, depending on the specific geometry at the scale of the obstacles (i.e. cells) [12]. In the absence of an exact scaling relationship for the cell geometry that we observe in experiments, we assume that the effective diffusivity for Nodal and Lefty follows a linear relationship (see also main text Fig. S3j)

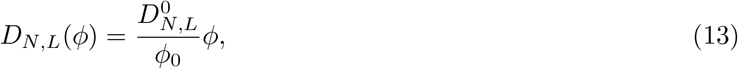

where 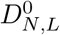 is the value of effective diffusivity at *ϕ* = *ϕ*_0_ (the initial value of porosity).

We now discuss how effective degradation and production terms scale with porosity. Consider *N*_*c*_ circular cells of radius *R* (see Figure SN1). The number of ligands internalised into the cells per unit of time is *k*_*int*_*N*_*c*_*σRc*_*ext*_, where *c*_*ext*_ is the extracellular concentration of ligands and internalisation happens with a rate *k*_*int*_ from a ring of width *σ* around the cell membrane. Using Eq. (11) this can be rewritten as

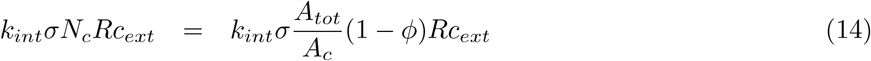

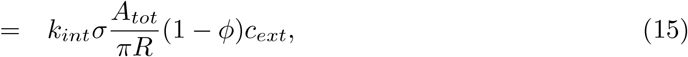

where in the last step we used *A*_*c*_ = *πR*^2^. We are seeking an expression in terms of the coarse-grained concentration *c*. At small porosities we can write Eq. (3) as *c* ≈ (*ϕ* + *β*)*c*_*ext*_. Then the total loss of ligands per unit of time can be written as

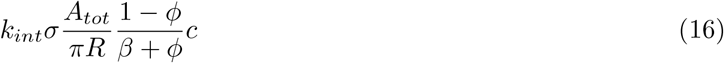

and one identifies the effective degradation as the prefactor of *c*, with the scaling

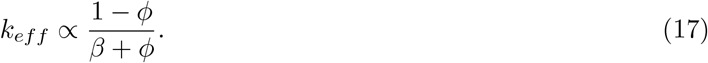

The effective degradation is thus an increasing function of the cell packing density. In the linear degradation terms in Eq. (6) and (7) we set

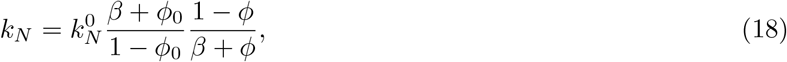

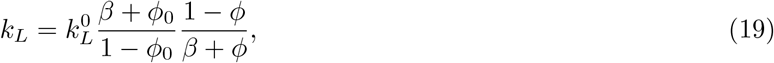

with 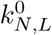 the values at *t* = 0. In the limit *β ≪* 1 and *ϕ ≪* 1, we obtain *k*_*eff*_ *∝* 1*/ϕ* and so effective degradation would increase inversely proportional to the cell distance [13].

We can obtain an expression for *β*, considering also degradation of internalised molecules with rate *k*_*o*_,

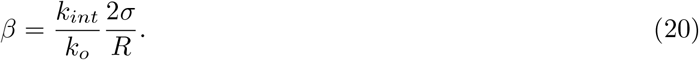

A similar scaling can be obtained for effective production. If one cell produces ν_*c*_ molecules per time, then in the tissue we will have *N*_*c*_ν_*c*_ molecules produced per time. Therefore the effective production per area is

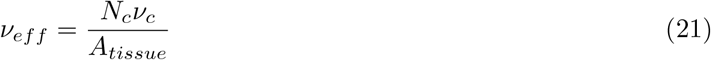

Since *A*_*tissue*_ = *N*_*c*_*A*_*c*_ + *A*_*ext*_, we can express *N*_*c*_ = (*A*_*tissue*_ − *A*_*ext*_)*/A*_*c*_ and

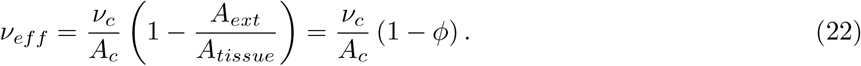

If the cell area *A*_*c*_ stays constant as *ϕ* changes, then the effective production scales as (1 − *ϕ*) with porosity. Thus, the higher the packing, the more molecules per area of tissue are produced. In equations (9) and (10) we set

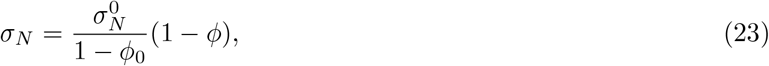

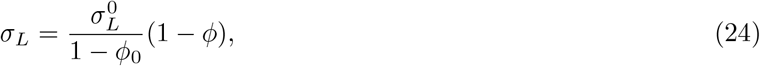

where 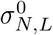 are the values set at *t* = 0.

### 1.4 Regulation of cell-cell adhesion and the relative surface tension *α* via Nodal

We now introduce the dynamics of the local porosity as a function of the Nodal concentration. As shown in the main text (Fig. 2a-c and Fig. S3d, d’) cell-cell adhesion strength is increasing with higher local Nodal levels. We introduce an effective, Nodal-mediated adhesion variable *E*, whose dynamics follows

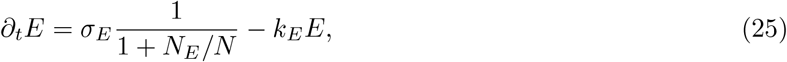

with an activation threshold *N*_*E*_ and degradation rate *k*_*E*_ that sets the time scale of adhesion build up. We set *E*(*x, t* = 0) = 0, such that this adhesion parameter captures only the additional adhesion strength beyond the baseline level at *t* = 0. Note that this is an effective parameter that captures local adhesion factors as a function of Nodal. For *N ≫ N*_*E*_, we would obtain the steady state value *E*^*\**^ = *σ*_*E*_*/k*_*E*_, which in turn is determined by the range of the relative surface tension *α* (see below).

We then describe the impact of this effective adhesion strength on *α* (introduced in the main text) as an algebraic relationship, such that *α* decreases with *E* as

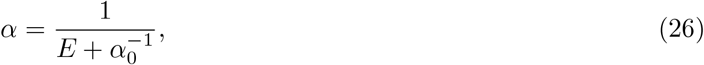

such that the simulation starts with the initial value of *α*(*x, t* = 0) = *α*_0_ = 0.9 for all positions *x* as measured in experiments (see main text, Fig 1g’). We set the lower bound to be *α*_*min*_ = 0.7, so that *α* varies in the experimentally relevant range (see main text, Fig. 1g’). This would be reached at maximal adhesion, and therefore sets 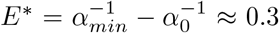 and *σ*_*E*_ = *E*^*\**^*k*_*E*_ = 0.3*k*_*E*_. In this way Eq. 26 is constrained by experimental measurements and does not introduce new free parameters in our model. Note that parameters that define *E* dynamics (i.e. *k*_*E*_ and *N*_*E*_) can then be tuned to modulate the *α* dynamics as a function of Nodal. Finally, *α* is updated in time irreversibly, i.e. at each position it can only decrease in time. This reflects the stability of adhesive contacts between cells.

### 1.5 Derivation of porosity as a function of *α*

To link tissue porosity to the relative surface tension *α*, we consider that the porosity of a tissue can be split into the individual contributions of the different gaps between cells, depending on the cells defining it. We have triangular gaps (surrounded by 3 cells), quadrilateral gaps (surrounded by 4 cells), pentagonal gaps (surrounded by 5 cells) and so on. Therefore, the porosity index *ϕ*(*α*), can be rewritten as:

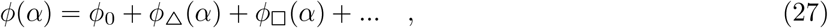

where *ϕ*_0_ is the minimum gap size, imposed by physiological constraints, and *ϕ*_△_(*α*), *ϕ*_□_(*α*) the contribution of each gap type as a function of *α*. To characterize the different terms, we will consider circles of radius *R* = 1. The center of masses of adjacent circles in the geometric arrangement defining the gap will be separated by distance *ℓ* = 2*R*. This distance will be kept constant throughout all the forthcoming developments. The area occupied by the cells will be computed by rescaling the radius of the circles –thereby formally triggering overlap– by an abstract scaling factor *ξ ≥* 1. The overlap will define an angle that can be related to the scaling parameter *ξ* and the relative surface tension parameter *α* derived in [14]:

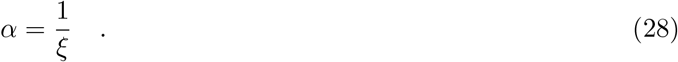

**Figure SN2:**
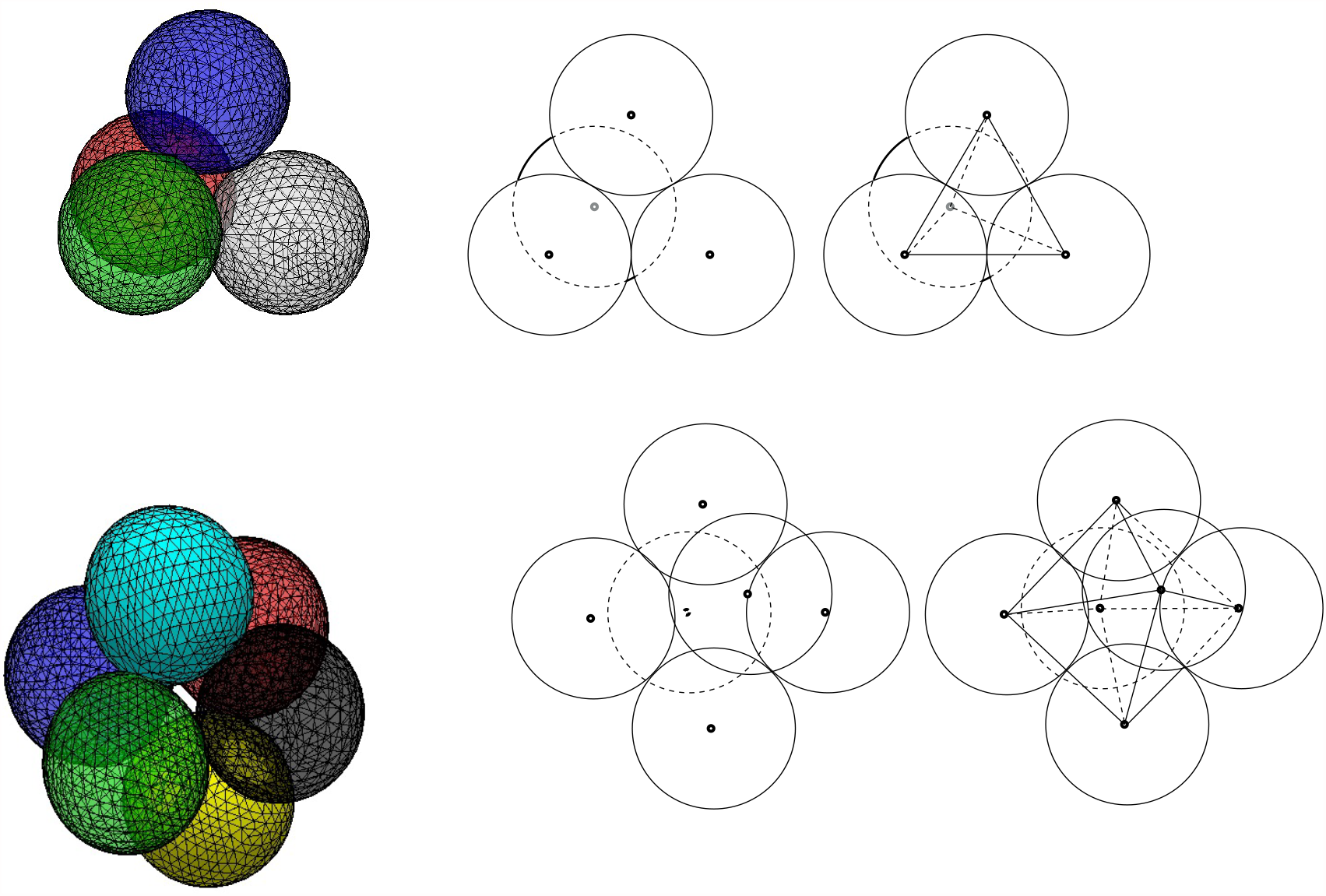
Packings in dense tissues –approximated by FCC crystallographic structures– show triangular, tetrahedral, octahedral and quadrilateral gaps. In 2D projections we consider only triangular and quadrilateral gaps.

We assume that the circles for *α* = 1 –hard disks– have radius *R* = 1 and *ξ* = 1. In the following section we will study the porosity from the analysis of the area covered by triangular and quadrilateral gaps. We consider these gaps because we approximate the structure of the 3D tissue to be close to the *face-centered cubic* (FCC) packing [15]. The projection of the FCC packing structure in 2D leaves triangular and quadrangular gaps –see Fig. (SN2) and Fig. (SN3).

#### 1.5.1 Triangular gaps

For each triangular gap, we consider the surface of triangle formed by the three centers of mass of three cells and how this is getting filled along the increase of *ξ*. Considering the fact that the triangle contains 1*/*6 of each of the 3 cells and the 1*/*2 of the intersection surface shared by each pair of cells, we have that the extracellular relative surface, *ϕ*, occupied by the cells within the triangle is:

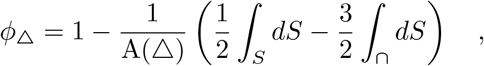

This leads to an equilateral triangle of side 2, being its overall surface of the 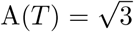. After applying a rescaling operation *R → ξR*, the other terms read:

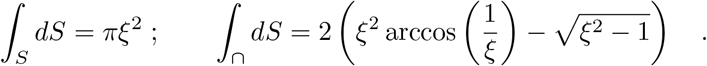

In terms of *α*, using that 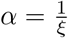:

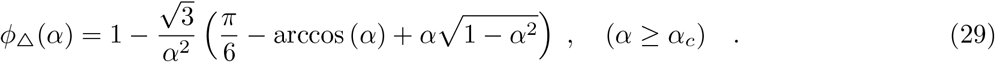

Triangular gaps are closed at 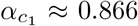. Even closed, there might remain some space open allowing the transport of morphogen molecules. This is grasped by the constant *ϕ*_0_. As we can observe in figure (SN3), even allowing little openings, the closing of triangular gaps at 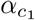 implies a collapse or drastic reduction of the connectivity of the interstitial network.

**Figure SN3:**
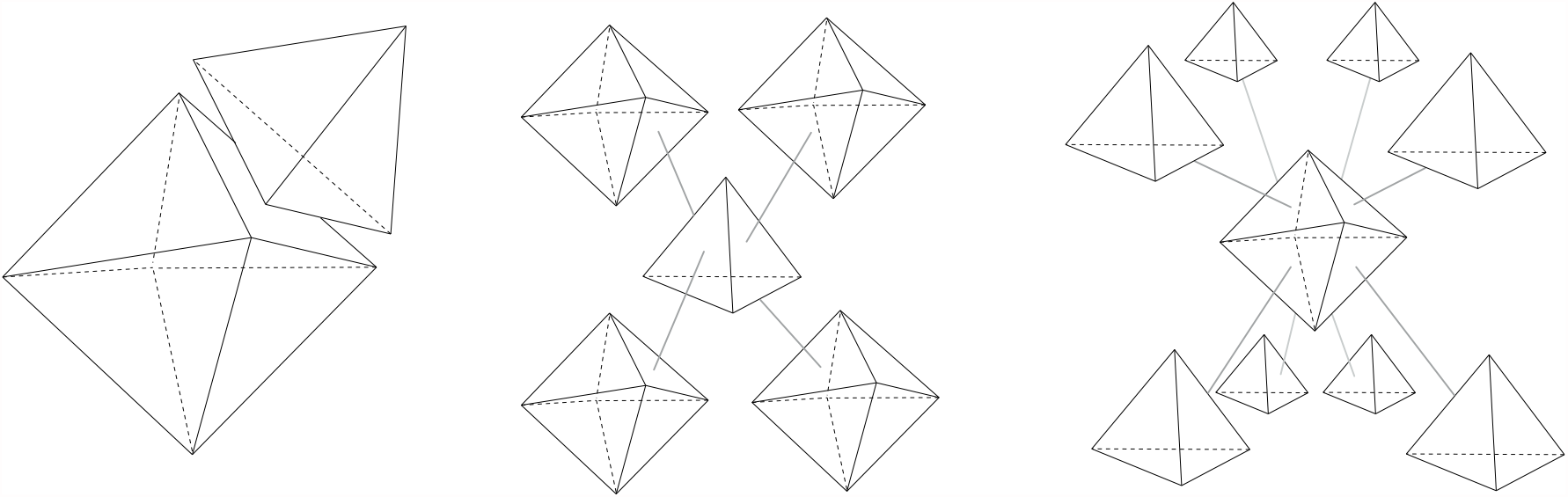
Local structure of the *gap network* of a Face-Centered Cubic (FCC) packing. Vertices describe the center of mass of the cells, connections describe cell-cell contacts. Interstitial channels cross the triangular faces. When they are closed at 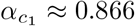 the whole network either collapses or, suffers a drastic reduction of its connectivity. However, octahedral structures, with a square-like configuration in the center may retain certain fluid that may disappear below 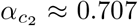, where, in force balance, no gaps are allowed any more in densely, FCC-like packings.

#### 1.5.2 Quadrilateral gaps

Using analogous reasoning, we can derive the relation accounting for the contribution of quadrilateral gaps. The general form will be:

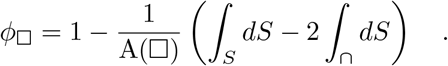

The area of the square will be *A*(□) = 4, leading to:

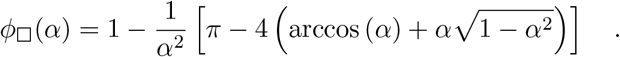

Using the relation described in equation (28), we can infer the *α* value at which quadrilateral gaps are closed, by just computing the re-scaling parameter *ξ* needed to reach the centroid of the square. A straightforward calculation leads to 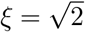. Therefore, we identify a second critical point below which, in a densely packed tissue, no interstitial gaps are expected, 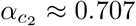. Using the same reasoning, we observe that the central point of a tetrahedron is reached at 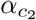. Consistently:

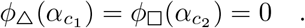

#### 1.5.3 Approximation for the porosity index

In a densely packed tissue, the main contributions are driven by square-like gaps and triangular-like gaps –see figures (SN2) and (SN3). We neglect the role of tetrahedral gaps, as they reveal vanishingly small in 2D projections. In these regards, comparison of equation (27) with real data shows a simple form:

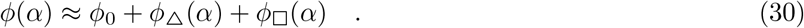

That is: just the sum of the triangular and quadrilateral contributions. We assume a contribution of *ϕ*_0_ = 10^−3^ for the minimum porosity attainable. This is consistent with experimental measurements in highly rigid and packed tissues (Fig. 3d in the main text). With this offset we take into account that diffusive transport, albeit slowed down considerably [16], never completely vanishes in a packed tissue. Expression (30) closes the system of equations and provides a feedback mechanism from Nodal signal, via *ϕ*(*α*), to the transport and kinetic parameters in the Nodal dynamics.

## 2 Numerical solutions

We solve equations (6), (7), (25) together with the relations (26) and (30) numerically on [0, *l*] with a uniform spatial grid with spacing *dx* = 1. Spatial derivatives are discretised and equations (6), (7), (25) are integrated in time using the Euler scheme with *dt* = 0.9(*dx*)^2^*/*(2 max {*D*_*N*_, *D*_*L*_}), according to the stability criterion for diffusion.

We begun by basing our analysis within the parameter ranges identified in [1]. Note, that there the parameter fits were performed using Nodal and Lefty profiles at the time of 50% epiboly (*t*=120min). Here, we instead consider both the intermediate dynamics and the steady state profiles.

We initially keep the diffusivities, degradation and production rates for Lefty and Nodal constant in time and space. In this case, the Nodal-Lefty system (Equations (6)-(7)) displays a non-monotonic Nodal dynamics (Fig. SN4 a), reminiscent of the one we find in experimental quantifications shown in the main text. At steady state, there is a Nodal gradient with a physiological range and a basically flat Lefty profile. At intermediate times, the Nodal gradient extends further into the tissue, but retracts as the amount of Lefty increases.

We then consider the full system of equations including the feedback through porosity, with parameters given in Table 1. We find that generally this enhances the non-monotonic dynamics of Nodal, with Nodal gradients retracting to a sharp profile (i.e. with reduced decay length) with higher levels of Nodal localised near the source (Fig. SN4 a versus SN4 b).

**Table 1:** Numerical values of parameters for Figure SN4.

| Parameter | Numerical value | Explanation |
| --- | --- | --- |
| $D_N^0$ | $1.95 \mu m^2/s$ | Nodal effective diffusivity at $t = 0$; Average of $0.7 \mu m^2/s$ (Cyclops) and $3.2 \mu m^2/s$ (Squint) from [4] |
| $D_L^0$ | $15.0 \mu m^2/s$ | Lefty effective diffusivity at $t = 0$; Average of $11.1 \mu m^2/s$ (Lefty1) and $18.9 \mu m^2/s$ (Lefty2) from [4] |
| $l$ | $300 \mu m$ | size of experimental measurement window |
| $k_N^0$ | $1.25 \times 10^{-6} s^{-1}$ | Nodal eff. degradation rate at $t = 0$ |
| $k_L^0$ | $7.5 \times 10^{-7} s^{-1}$ | Lefty eff. degradation rate at $t = 0$ |
| $k_{NL}$ | $10 s^{-1}$ | Lefty-Nodal inhibition parameter; range $10^{-5} - 10^{-1}$ in [1] |
| $N_a$ | 31.62 a.u. | Nodal production activation threshold |
| $N_L$ | 300 a.u. | Lefty production activation threshold |
| $L_N$ | 15.8 a.u. | Lefty-induced inhibition threshold |
| $\sigma_N$ | $10^{-2}$ a.u. | Nodal (relay) production strength |
| $\sigma_L$ | $1.2 \times 10^{-2}$ a.u. | Lefty production strength |
| $k_E$ | $2 \times 10^{-4}$ | sets timescale of adhesion strength build-up |
| $N_E$ | 23.71 | $0.75 \times N_a$ |
| $\sigma_E$ | $5.88 \times 10^{-5}$ | $\sigma_E = E^* k_E$ |
| $E^*$ | 0.2941 | $E^* = \alpha_{min}^{-1} - \alpha_0^{-1}$ |
| $\alpha_0$ | 0.881 | $\langle \alpha(x) \rangle (t = 0)$ from experiment (WT, see main text) |
| $\alpha_{min}$ | 0.7 | minimal $\alpha$ from experiment |
| $\beta$ | 0.1 | $\beta = c_{int}/c_{ext}$ |
| $m_N$ | 2 | Hill-coefficient in Nodal relay |
| $m_L$ | 8 | Hill-coefficient in Lefty production |

To quantify the spatiotemporal dynamics of the Nodal range we follow the method outlined below (see Fig. SN4 b):

- Define *λ*_1*/*2_(*t*) as the distance at which the concentration reaches *N*(*x* = 0, *t*)*/*2 for each time *t*. This is a dynamical variable that captures the range of the Nodal gradients over time.
- Find the time *τ* = argmax_*t*_ *λ*_1*/*2_(*t*). This time point indicate the time at which the nodal gradient is the widest and so the time point just before the Nodal gradient begins to retract.
- Define *N*_*max*_ = *N*(*x* = 0, *τ*), that is the maximum Nodal concentration at the time point when the Nodal range is maximised. We then quantify the range of the Nodal profile over time as the position *x*_*N*_ at which the gradient reaches the *p* fraction of that value, i.e. *N*(*x*_*N*_) *< pN*_*max*_, with *p* = 0.1, 0.2, 0.3.
- We normalize time by defining *τ*_*WT*_ to be the time of the peak in the Nodal length scale profile and setting *τ*_*WT*_ = 1.

This approach allows us to assess the position beyond which the Nodal signalling levels drop below a pre-defined threshold. This method of quantifying the Nodal range is designed to be comparable to the experimental quantifications in the main text where we quantify the Nodal range by measuring the number and extend of Smad2 positive nuclei, presumably only detecting nuclei that have sufficiently high levels of Smad2. The quantifications presented in Fig. 3 in the main text take *p* = 0.2. Note that the value of *N*_*max*_ is quantified from the WT numerical solutions (with feedback) and the same value of *N*_*max*_ is used to define the lower threshold for signal detection in WT and wnt11 solutions (with and without feedback respectively).

### 2.1 Quantifications

We now compare simulations that represent the following experimental conditions: WT (full model; labelled *with feedback*), MZwnt11/slb mutant (*E, α, ϕ* dynamics turned off: *σ*_*E*_ = 0; labelled *without feedback*), MZlefty1/2 mutant (*L* production turned off: *σ*_*L*_ = 0; labelled *without lefty*).

Throughout our analyses we compare model dynamics to the experimental observations by quantifying the following measures:

- the Nodal range as defined above, with *p* = 0.2
- time *τ*_*WT*_ of Nodal range peak in the WT simulation and normalise time *t* relative to this
- the average porosity 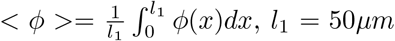, corresponding to the IFF measurement close to the YSL.
- the total production of Lefty and Nodal in the tissue over time: 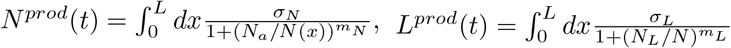.
- spatiotemporal profiles of Nodal *N*(*x, t*), which we compare to the experimental profiles of Smad2 N/C ratios

**Figure SN4:**
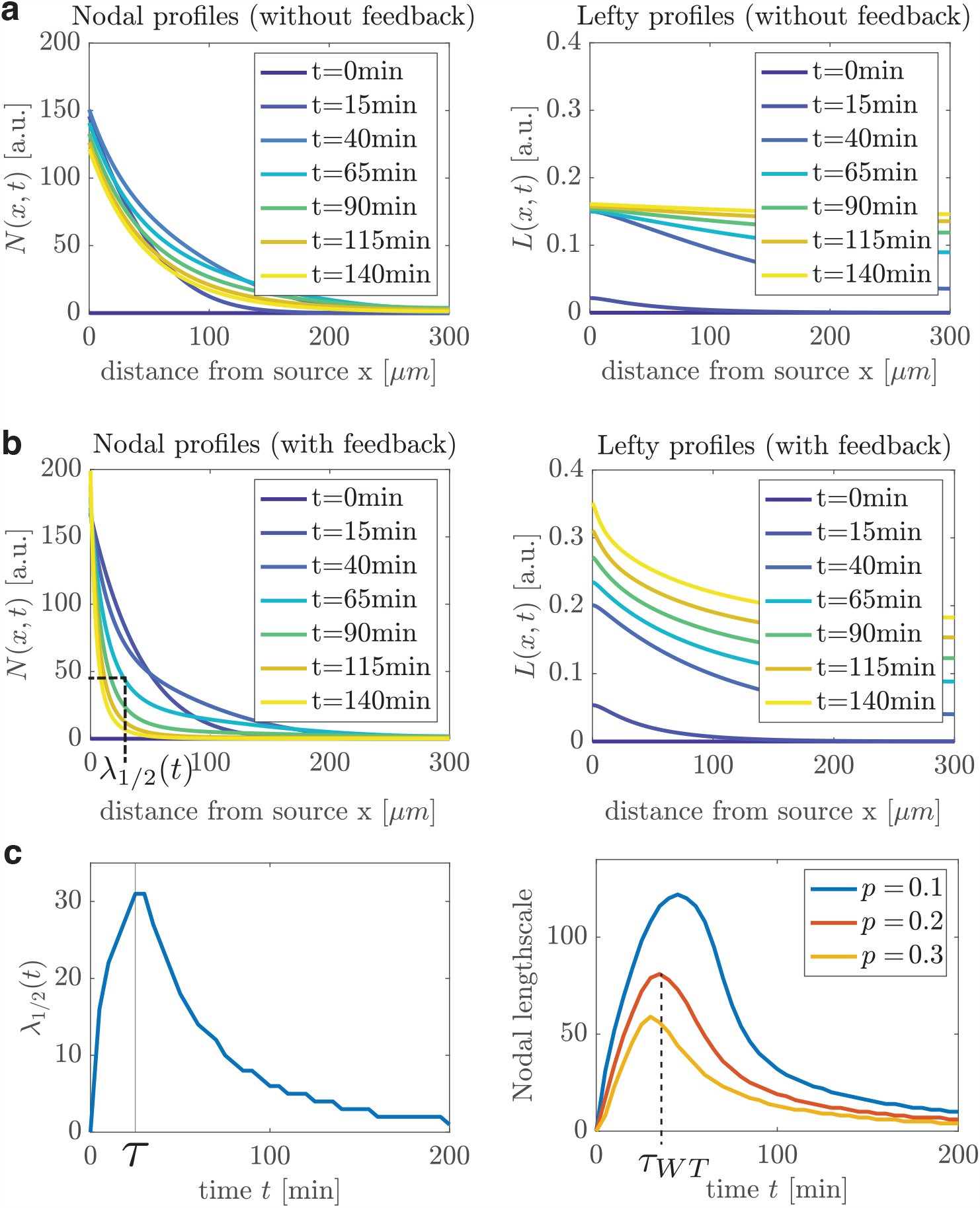
**(a)** Dynamics of Nodal and Lefty with constant diffusivities and degradation and production rates (i.e. without feedback). **(b)** Dynamics of Nodal and Lefty with feedback through porosity. c) Quantification of the non-monotonic dynamics of the Nodal range, with the time of Nodal peak *τ*_*WT*_ defined on the *p* = 0.2 curve. Parameters are as given in table 1, but with *σ*_*E*_ = 0 (i.e. porosity dynamics is turned off).

### 2.2 Parameter choices

Table 2 shows the parameters used for the theoretical model results shown in figures 3 and 4 in the main text. Although the expansion of the Nodal gradient and a delay in Nodal termination appear to be general features of the theoretical set-up with feedback between porosity and Nodal, we have identified some dependencies of these behaviours on parameter choices.

**Table 2:** Numerical values of parameters for main text Figures 3 and 4 (parameters not listed here are the same as in Table 1).

| Parameter | Numerical value |
| --- | --- |
| $k_N^0$ | $1.5195 \times 10^{-6} s^{-1}$ |
| $k_L^0$ | $7.4653 \times 10^{-7} s^{-1}$ |
| $k_{NL}$ | $10.3034 s^{-1}$ |
| $N_a$ | 31.6228 a.u. |
| $N_L$ | 200 a.u. |
| $L_N$ | 19.1507 a.u. |
| $\sigma_N$ | $10^{-2}$ a.u. |
| $\sigma_L$ | $4.6539 \times 10^{-4}$ a.u. |
| $k_E$ | $2.3082 \times 10^{-4}$ |

The observed features depend on the following relations between the activation thresholds for the three key model components that depend on the local Nodal levels: *N*_*E*_ *< N*_*a*_ *< N*_*L*_. In other words, as the Nodal profile grows, the following temporal sequence drives the dynamics we observe

- first, the adhesion strength begins to increase, leading to a drop in the porosity near the margin
- second, the Nodal relay is activated, extending the Nodal gradient further into the tissue
- third, Lefty levels start to rise and eventually cause the Nodal profile to retract. The porosity-mediated localisation of both Nodal and Lefty near the source occur on a similar time scale, in order to enhance the inhibiting effect of Lefty.

For the Lefty production to be responsive to the porosity-induced changes of Nodal dynamics, the activation threshold for Lefty, *N*_*L*_, needs to lie approximately at the maximum Nodal values achieved in the no-feedback (MZwnt11/slb) simulations. In this way, the porosity-mediated localisation of Nodal at the source enhances local Lefty production. This, in turn, reinforces the biochemical feedback between Lefty and Nodal (as shown by L-N degradation term).

Further, parameters are chosen to ensure that in the absence of Lefty, the Nodal gradient extends throughout the entire tissue (see Fig. SN5), i.e. the porosity change alone is not sufficient to cause Nodal retraction. Imposing this constraint in our theoretical model is motivated by our experimental quantifications of the Smad2 positive nuclei in Lefty mutants, where Smad2 positive nuclei extend to the entire tissue (Fig. 2e in the main text).

To improve our parameter estimation we used the Mesh Adaptive Direct Search algorithm (MADS), designed for difficult blackbox optimization problems [17]. These issues occur when the functions defining the objective are the result of costly computer simulations, which in this work arise since we are solving the dynamics of Eq.(6), (7), (25) numerically. The loss function was defined as the sum of two contributions. First, the mean square error between the experimentally measured Smad2 positive nuclei distance from the YSL over time in WT and MZwnt11/slb conditions (Fig. 3g and h in the main text) and the corresponding range of the Nodal signalling gradient with and without feedback that we computed as outlined in the previous subsection. Second, the loss function accounts for the mean square error between the *α* quantifications in space and time in the WT data and the corresponding values of *α* computed numerically; Fig. S3e cited in the main text shows experimental data and best theoretical fit found by the optimizer for *α*(*x, t*). These two mean squared errors were individually normalized so that their contributions were equal. This was run locally to the parameter sets we had identified from previously published work and using the analysis above.

**Figure SN5:**
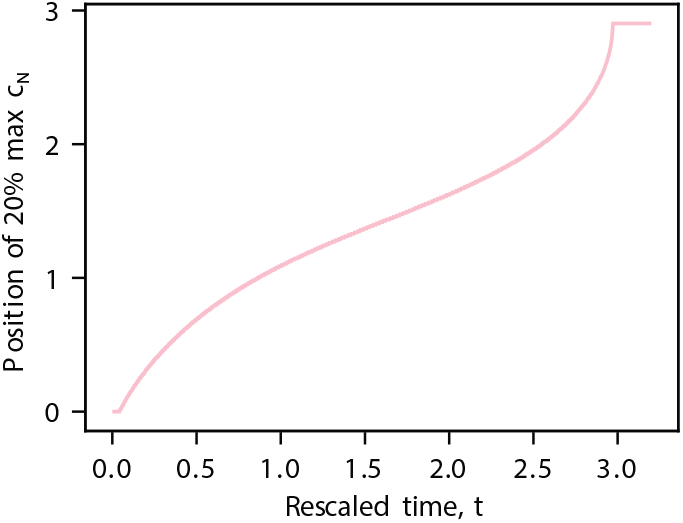
The range of Nodal length scale in numerical solutions without Lefty (*σ*_*L*_ = 0) for the set of parameters used to produce all main text figures. The max *C*_*N*_ used here is based on the case with feedback in Fig. 3f (main text). Parameters: see Table 2. The length scale reaches saturation, i.e. Nodal extends through the entire tissue.

### 2.3 The role of Nodal relay

Nodal has been proposed to disperse through diffusion of ligands secreted from the YSL and a relay mechanism whereby Nodal signalling activates its own transcription and secretion [5]. We have therefore implemented both of these processes to our theoretical framework and analysis.

Previous work shows that a relay mechanism can lead to production in the entire tissue in certain parameter ranges [18]. To estimate the relay strength *σ*_*N*_ for which this would happen in our model and define parameter regimes where relay operates, we consider the Nodal equation with *m*_*a*_ = 1, in the absence of Lefty and the external source,

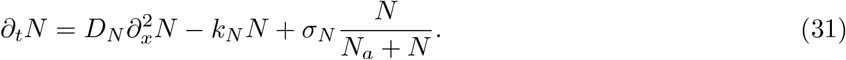

The steady state equation has two constant solutions

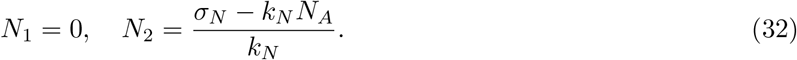

However, *N*_2_ *>* 0 (i.e. the non-trivial solution exists) only if *σ*_*N*_ *> k*_*N*_ *N*_*a*_, which provides an estimate for the critical activation strength beyond which the entire tissue will be producing, also in the case of a non-zero source.

To investigate whether our findings hold true, independently of the relative contribution of diffusive versus relay-driven mechanisms, we have varied the contribution of the two processes and asked whether the key features we have identified in our analysis in the presence of feedback between Nodal and tissue packing are general. We found that the reduction in the Nodal range and the faster termination of Nodal signalling we report in the main text are consistent with parameter choices that remove the relay contribution all together and that enhance the relay contribution compared to the parameter set used for our main text results (Fig. SN6a). In both these cases we noted a delay in lefty production in the absence of the feedback (Fig. SN6b) and expansion of the Nodal gradient and reduction in the Nodal concentration near the margin (Fig. SN7), as reported in the main text. To estimate the contribution of the relay mechanism to the total gradient we compared *λ*_1*/*2_(*t*) for the case with relay (*σ*_*N*_ = 0) and the case where we turn relay off (*σ*_*N*_ = 0) (Figure SN6, c).

**Figure SN6:**
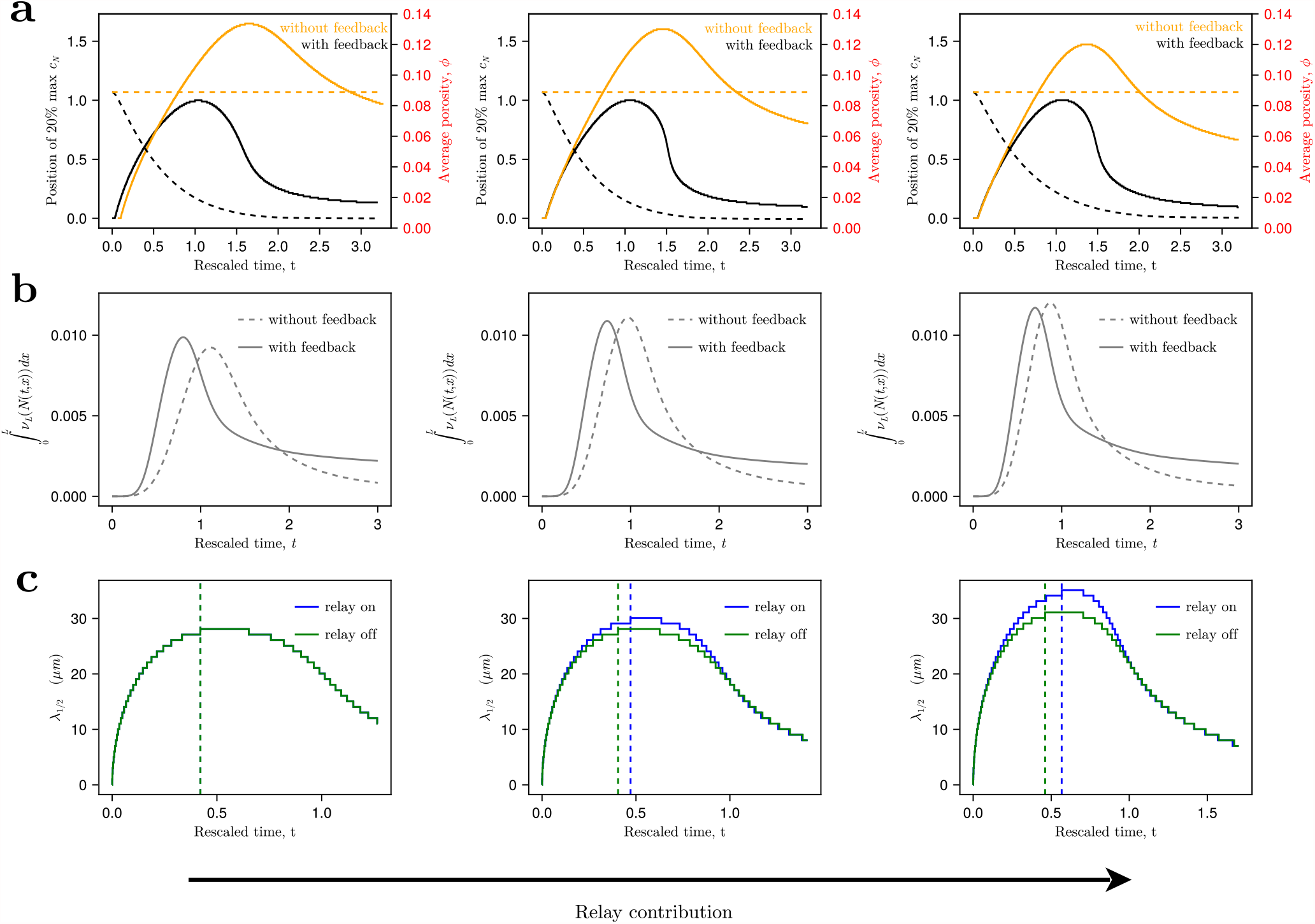
**(a)** Nodal length scales and averaged porosities from simulations with and without feedback for varying contribution of the Nodal relay component as indicated by the arrow at the bottom of the figure. **(b)** Total Lefty production in the tissue with and without feedback. **(c)** *λ*_1*/*2_(*t*) from simulation with feedback, with and without relay. Parameters: for left column as in main text Fig. 3, 4 but *σ*_*N*_ = 0, for middle column as in main text Fig. 3, 4, for right column as in main text Fig. 3, 4 but 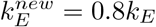 and 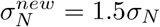.

**Figure SN7:**
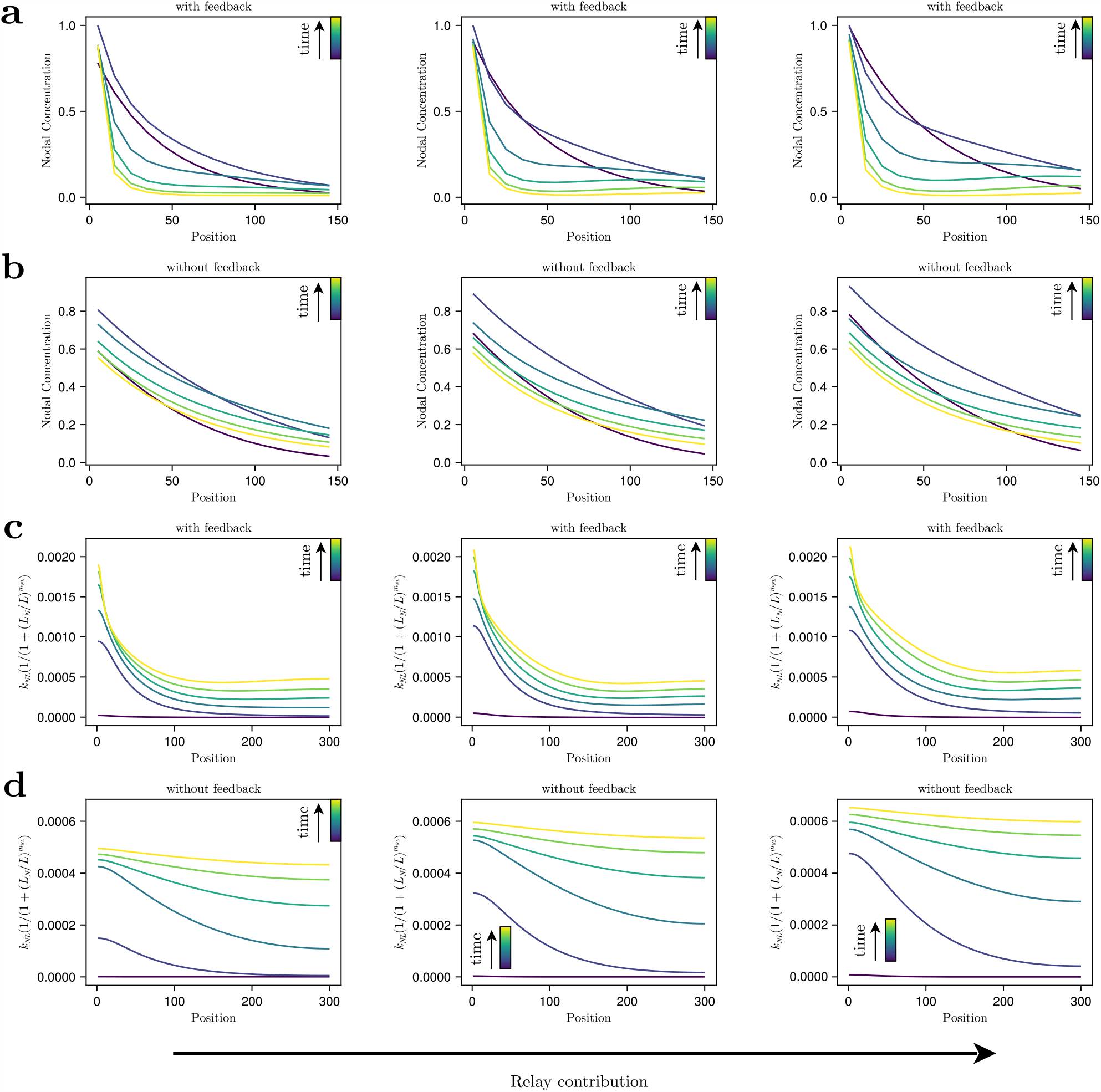
Nodal concentration profiles with feedback **(a)** and without feedback **(b)** (both scaled to maximum value in (a)), and magnitude of Lefty-induced inhibition with feedback **(c)** and without feedback **(d)**. Time points plotted are [0.5, 1, 1.5, 2, 2.5, 3]*τ*_*WT*_ (from dark blue to yellow). Columns map to parameters that correspond to an increase contribution of relay from left to right as indicated by the arrow at the bottom of the figure. Parameters: for left column as in main text Fig. 3, 4 but *σ*_*N*_ = 0, for middle column as in main text Fig. 3, 4, for right column as in main text Fig. 3, 4 but 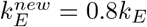 and 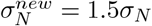.

## 3 Impact of diffusivity and degradation on the morphogen amplitude and decay length

To understand how changes in the effective diffusion, *D*_*eff*_ and effective degradation *k*_*eff*_ are expected to affect the morphogen amplitude (*C*_0_) and the decay length *λ*, we consider the minimal diffusion-degradation equation for a morphogen [19]

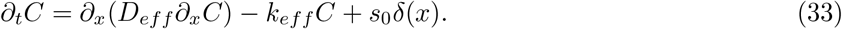

with no-flux boundary conditions *∂*_*x*_*C*|_*x*=0_ = *∂*_*x*_*C*|_*x*=*l*_ = 0. In the limit *λ ≪ l*, the (well-known) steady state is

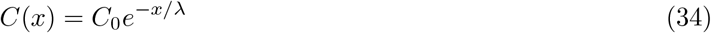

with the decay length 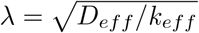 and amplitude 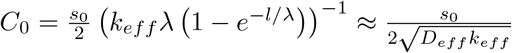. Figure SN8 illustrates that, in this simple scenario, an increase in *C*_0_ can be facilitated by a reduction in *D*. This reasoning indicates that the changes in the GFP tagged Squint observed experimentally are consistent with the conclusion that the tissue rigidity transition and concomitant decrease in porosity restricts the diffusivity of Nodal ligands (main text Fig. 3m-o and Fig. S3i-l).

**Figure SN8:**
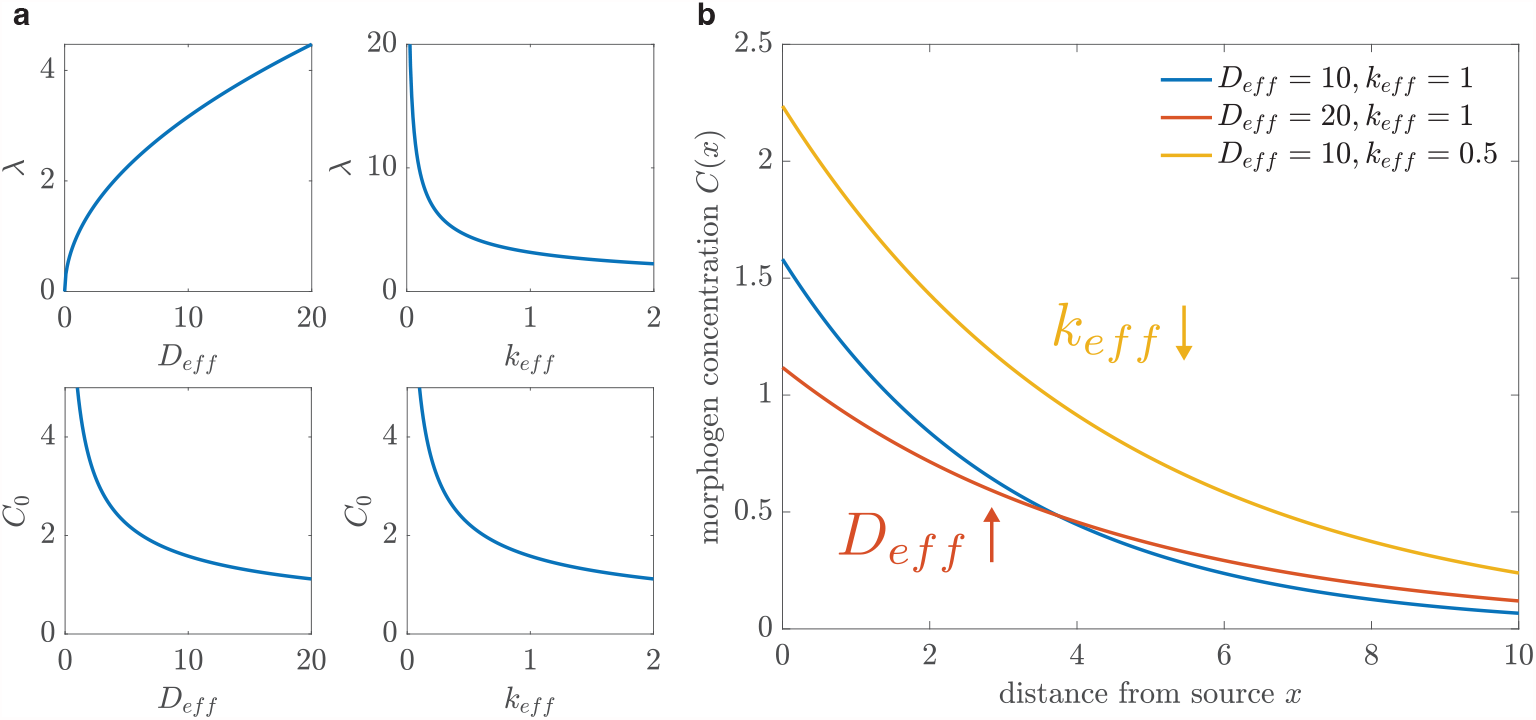
**(a)** Plots of 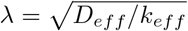 and 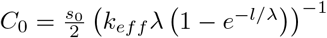. **(b)** Halving the degradation or doubling the diffusivity has the same effect on *λ*, but the amplitude increases in the former case and decreases in the latter. Parameters (if not varied): *s*_0_ = 10, *l* = 100, *D*_*eff*_ = 10, *k*_*eff*_ = 1.

## 4 Simulations of cell networks and cell tilings with varying connectivity and adhesion

In this section we detail the steps followed to construct networks and cell tilings with gradient in adhesion and connectivity along the vertical axis. For the latter, we introduce the soap-bubble Hamiltonian [20, 21], which will become the target function that will lead the optimization process. From the structure of cell-cell contacts, we can derive the rigidity properties of the system. Besides the emerging topological structure of the network of cell-cell contacts, the computation of optimal tilings given a certain surface tension parameter is also used to compute the formation of Tri-cellular junctions.

### 4.1 Construction of networks with connectivity gradient

The basic network structure is a 2D triangular lattice, in which some disorder in the location of nodes is introduced, which may trigger the appearance of new links, due proximity, or their removal, in case the two nodes get separated more than a threshold beyond which it is considered that they cannot represent a cell-cell contact. Upon this basic structure, to explore how the cell connectivity affects the spatial distribution of the GRC, we generated networks of size 35×35 nodes with a linear connectivity gradient, keeping the global average connectivity fixed. This is achieved by pruning links probabilistically from a totally connected triangular lattice –with geometric disorder, as explained above– with link deletion probability increasing linearly along the vertical axis of the network.

### 4.2 Construction of in-silico cell-tilings with adhesion gradient

To explore the effects of the adhesion gradient on the GRC localization, we simulate non-confluent tissues as 2D cell arrangements with linearly decreasing adhesion along the vertical axis. The energy of the tissue can be described by a soap-bubble-like Hamiltonian [20, 21]:

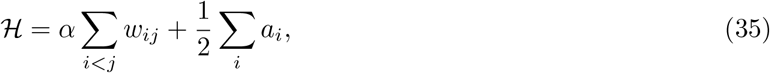

where *w*_*ij*_ denotes the contact area between cells *i* and *j, a*_*i*_ is the area of cell *i* in contact with interstitial fluid, and *α* is a non-dimensional parameter defined as:

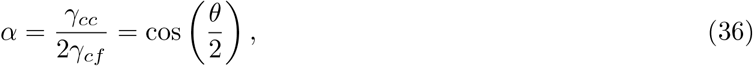

with *θ* being the angle between the membranes of two cells in contact, *γ*_*cc*_ is the cell-cell surface tension, and *γ*_*cf*_ is the cell-fluid surface tension. The above equation establishes a connection between the relation among surface tensions and the angle between two membranes in contact to the fluid, an observable feasible to extract from real systems. Numerical simulations of 2D cell tilings have been performed using the C based software Surface Evolver version 2.70 [22]. To start the simulation, the initial configuration needs to be generated, specifying the location of the cells, a primary polygon-like geometry and the initial topology of cell-cell contacts. Later this initial condition will evolve 1/ By increasing the resolution of the perimeter of the cells –thereby achieving realistic geometries– and 2/ Optimizing the global geometry of individual cells and size of cell-cell contacts according to the global soap-bubble-like Hamiltonian as the one in Eq. 35, in a relaxation process towards the desired *α* value. The action of external pulling - or pushing forces is not considered here and, thereby, we expect the tilings to be in equilibrium with respect the soap-bubble Hamiltonian. To generate random tilings with subcritical target densities *ϕ*_*T*_, we follow the RSA algorithm [23] as implemented in [14] –further we will introduce the gradient in adhesion:

- Over an a-priori defined square of length *L* –in units of cell diameter *D* = 2*R*, where *R* is the average cell radius– send *N* ^*′*^ random possible coordinates 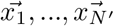.
- The sequential process of generation of random coordinates is subject to a selection criteria: If when generating the *k*-th random coordinate an already existing coordinate 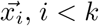 is such that 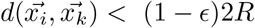 this coordinate 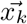 is discarded, as it would lead to a large overlapping pair of disks.
- If the number of accepted generated coordinates *N* (*N ≤ N* ^*′*^) reaches a value such that, *ϕ*_*T*_ *≤ NϕR*^2^/*L*^2^, where *ϕ*_*T*_ is the target density, the process stops, since the target density has been achieved.
- If after a long number of iterations, the target density cannot be achieved, we perform a random search along the area identifying possible empty spaces that can be filled using the previous distance conditions.
- We build a collection *C*_1_, …*C*_*N*_ of disks centered on each of the accepted coordinates 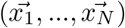. For any pair of disks *C*_*i*_, *C*_*k*_ such that 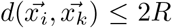 we define a contact. Since the presence of overlaps and further changes on adhesion may slightly alter the cell fraction, we may encounter the situation by which the actual achieved density *ϕ < ϕ*_*T*_. If this happens, we need to generate more disks, refining the halting condition and rewriting it as *ϕ ≤* (*N* + *δ*)*ϕR*^2^/*L*^2^ to achieve the desired densities *ϕ* ≈ *ϕ*_*T*_ after optimization. Once generated, the seed is stabilized in the hard-disk regime (*α* = 1). Each cell is described by a set of vertices and segments, whose minimum length is controlled by the parameter *t*. We generate tilings of 26×26 cells. The evolution of the tissue will be simulated as follows.
- Decrease *α* for each region of the tiling, according to the imposed gradient, in a quasi-static way until the desired *α* is reached. The gradient is imposed such that the bottom layer has target *α* = 0.77 and the top layer has target *α* = 0.95, values that agree with the measurements in real systems. In this step, the mechanism of re-meshing implemented by the `Surface Evolver` acts by alternatively reducing and increasing the resolution of the geometry of the cells. This is done by merging all segments lower than a certain length (*t* parameter) and subsequently dividing such segments by 2. With this, the optimization algorithm can explore the potential configurations minimizing the global surface energy in an efficient way. According to our simulations, working with very fine grained geometries (low *t* parameters) all the time may trap the system in local optima. In consequence, we need to introduce sporadically large merging events (large *t*’s) to trigger topological changes. To preserve the high resolution of the simulation but, at the same time, allow topological rearrangements, we apply the re-meshing *t*-parameter as following a Weibull function:

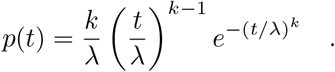 This allows us to perform most of the re-meshing events around a well defined mean, keeping high resolution, but, from time to time, massive re-meshing events allow topological rearrangements.
- When the targeted *α* is reached and no changes on the energy are appreciated throughout successive rounds of optimization, we end the process. The resulting configuration is a tiling with the desired *α* gradient and approximately the target density *ϕ*_*T*_.

### 4.3 Rigid cluster analysis

Identification of floppy and rigid areas of the 2D cell-cell contact networks was performed using pebble.py, available at: https://github.com/coldlaugh/pebble-game-algorithm/blob/master/pebble.pyx [24].

